# A Large Field-of-view, Single-cell-resolution Two- and Three-Photon Microscope for Deep Imaging

**DOI:** 10.1101/2023.11.14.566970

**Authors:** Aaron T. Mok, Tianyu Wang, Shitong Zhao, Kristine E. Kolkman, Danni Wu, Dimitre G Ouzounov, Changwoo Seo, Chunyan Wu, Joseph R. Fetcho, Chris Xu

## Abstract

In vivo imaging of large-scale neuron activity plays a pivotal role in unraveling the function of the brain’s network. Multiphoton microscopy, a powerful tool for deep-tissue imaging, has received sustained interest in advancing its speed, field of view and imaging depth. However, to avoid thermal damage in scattering biological tissue, field of view decreases exponentially as imaging depth increases. We present a suite of innovations to overcome constraints on the field of view in three-photon microscopy and to perform deep imaging that is inaccessible to two-photon microscopy. These innovations enable us to image neuronal activities in a ∼3.5-mm diameter field-of-view at 4 Hz with single-cell resolution and in the deepest cortical layer of mouse brains. We further demonstrate simultaneous large field-of-view two-photon and three-photon imaging, subcortical imaging in the mouse brain, and whole-brain imaging in adult zebrafish. The demonstrated techniques can be integrated into any multiphoton microscope for large-field-of-view and system-level neural circuit research.

## Introduction

High spatial and temporal resolution imaging over a large field-of-view (FOV) deep within intact tissues is valuable for many biological fields such as neuroscience. In particular, high-resolution imaging is essential to minimize the risk of misattributing calcium transients (***Gauthier et al., 2022***) in cellular calcium imaging. While large FOV two-photon microscopy (2PM) (***Sofroniew et al., 2016***; ***Lu et al., 2020***; ***Demas et al., 2021***; ***Stirman et al., 2016***; ***Yu et al., 2024***; ***Clough et al., 2021***; ***Ota et al., 2021***; ***Rumyantsev et al., 2020***; ***Tsai et al., 2015***; ***Janiak et al., 2022***) has demonstrated recording neural activities up to 5-mm FOV, the penetration depth of two-photon (2P) imaging is restricted to the shallow cortical layers in the intact mouse brain (***Wang and Xu, 2020***; ***Wang et al., 2020***; ***Takasaki et al., 2020***). Moreover, achieving large FOV activity imaging often comes at the expense of reduced imaging resolution. In contrast, three-photon microscopy (3PM) has reliably imaged neurons in deep cortical layers (***Takasaki et al., 2020***), subplates (***Yildirim et al., 2019***), and subcortex (***Weisenburger et al., 2019***; ***Ouzounov et al., 2017***), offering optical access to regions inaccessible to 2PM. The combination of 2PM and 3PM has enabled deep neural imaging in the neocortex and the sub-cortical region simultaneously with a limited FOV (***Weisenburger et al., 2019***) of ∼0.5 x 0.5 *mm*^2^. Recent advancement in 3PM has shown vasculature imaging up to 716 *μ*m in the intact mouse brain with 2-mm FOV (***Yu et al., 2024***). However, 3PM requires low laser repetition rates for deep imaging to avoid thermal damage. Since the maximum number of pixels per second is limited by the laser repetition rate (***Weisenburger et al., 2019***), 3PM for deep, single-cell-resolution neural activity imaging is constrained to a small FOV of hundreds of micrometers (***Wang and Xu, 2020***).

To address the limitations of 3PM, we have developed a Dual Excitation with adaptive Excitation Polygon-scanning multiphoton microscope (DEEPscope). DEEPscope has a large FOV and is capable of simultaneous 2P and three-photon (3P) deep neural activity imaging with single-cell resolution. We introduced a suite of innovations, including adaptive excitation to reduce the average power required for 3PM, point-spread-function optimization for higher excitation efficiency, and a scalable multi-focus polygon scanning method to achieve scan rate beyond the limit of a mechanical scanner. This work extends the FOV of 3PM from 0.25 *mm*^2^ to 9.6 *mm*^2^ to image the deepest cortical layer at single-cell resolution at 4Hz, overcoming previous constraints on 3PM FOV.

We demonstrated the performance of DEEPscope in the mouse cortex, hippocampus and adult zebrafish. We achieved large FOV three-photon structural imaging of neurons in an intact mouse brain to >1.1-mm deep, reaching the hippocampus, and simultaneous 3P and 2P recordings of more than 4,500 neurons in both the deep and shallow cortex. Our approach allowed for neuronal recording in an intact mouse brain with a 3.23 mm x 3.23 mm FOV in cortical layer 6, and a 550-*μ*m FOV in the mouse hippocampus with single-cell resolution. We performed high spatial-resolution 2P activity imaging with a 0.7-*μ*m pixel size at 4 Hz with a large FOV of ∼3.23 mm x 1 mm, where labeled neurons and dendrites are clearly visible. We also performed whole-brain structural imaging of adult zebrafish brain, with deep penetration of >1 mm.

The DEEPscope is modular, scalable, and readily integrated into any multiphoton microscope to improve its imaging throughput, and it offers a promising path towards achieving the full potential of multiphoton microscopy for high-throughput and high-resolution imaging of a large population of neurons deep and across a large FOV. Furthermore, DEEPscope is simple and compact, which will enable straightforward implementation in neuroscience laboratories.

## Results

### DEEPscope for large-field-of-view imaging deep in scattering tissue

We have developed DEEPscope that enables single-cell-resolution imaging with a large FOV (3.5 mm diameter) deep in scattering tissue. ***Figure 1*** shows the experimental setup. The microscope allows simultaneous 2P and 3P excitation (i.e., dual excitation). The 3P excitation path consists of an adaptive excitation module, a beamlet generation delay line, and a scan engine with a polygon scanner. The 2P excitation path consists of an adaptive excitation module, a remote focusing module, and the same polygon scan engine for 3P excitation. The DEEPscope achieved deep and large FOV imaging in scattering brain tissues by (1) performing fast and large-angle optical scans using the polygon scanner, (2) reducing the power required for large-FOV imaging using adaptive excitation, and (3) improving the excitation efficiency and scanning speed using optimized point spread function (PSF) and beamlets.

**Figure 1.**
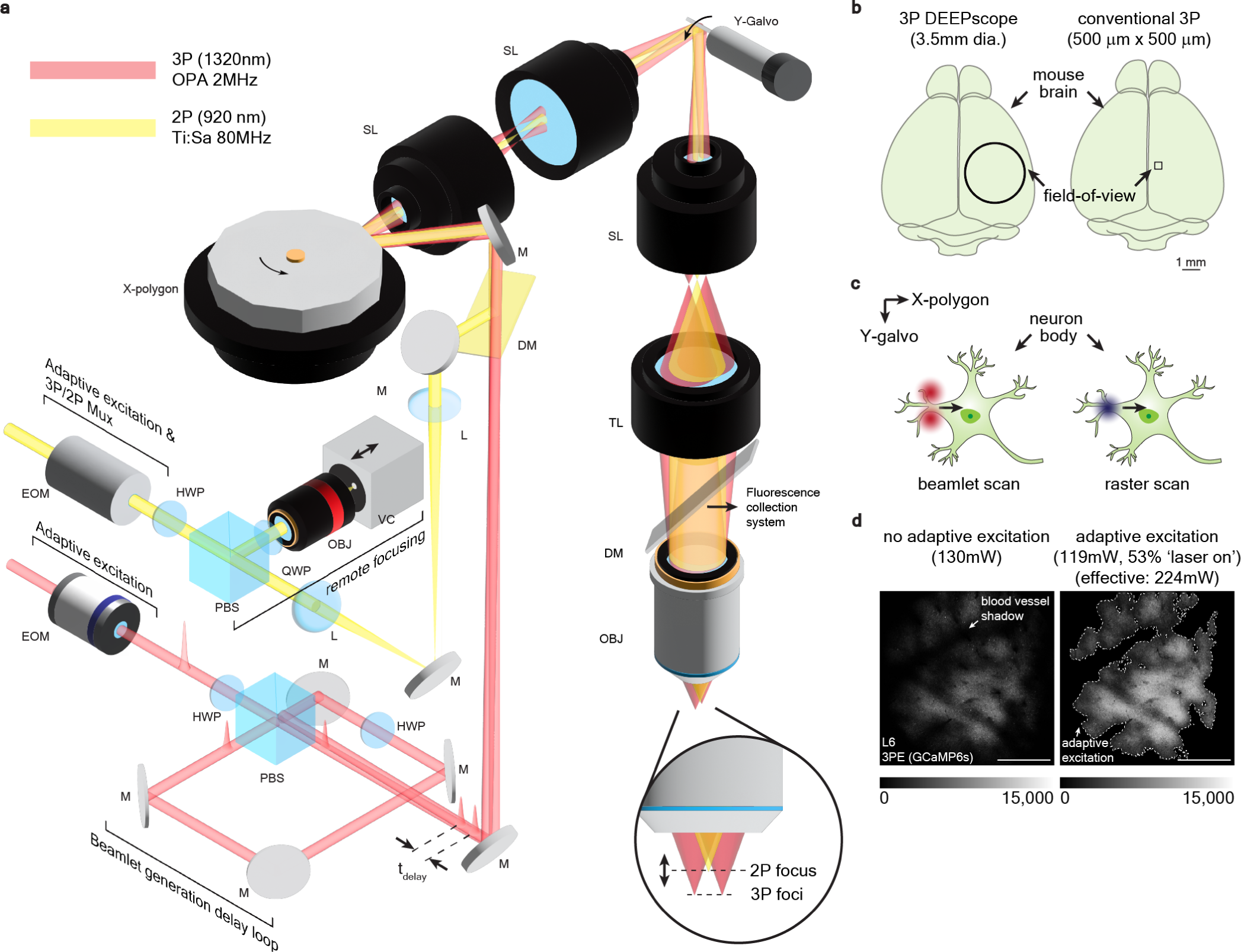
Dual-beam Excitation with adaptive Excitation Polygon-scanning multiphoton microscope (DEEPscope). **(a)** DEEPscope schematics. The 3P path consists of an adaptive excitation module, a beamlet generation delay loop, and a scan engine with a polygon scanner. The 2P path consists of an adaptive excitation module, a 3P/2P multiplexing module, a remote focusing module, and the same scan engine as for 3P imaging. **(b)** Comparison of the field-of-view (FOV) of 3P DEEPscope with conventional 3P microscopes. DEEPscope images 3.5 mm diameter FOV in the mouse brain. **(c)** Schematic of beamlet scanning with the beamlet generation delay loop. **(d)** Comparison of fluorescence intensity across the FOV with and without adaptive excitation at similar average excitation power. Grayscale bar shows pixel intensity. Scale bar, 1 mm. EOM: electro-optic modulator, HWP: half-wave plate, QWP: quarter wave-plate, PBS: polarization beam splitter, L: lens, M: mirror, SL: scan lens, TL: tube lens, DM: dichroic mirror, OBJ: objective, tdelay: time delay between the pulses of the two beamlets.

While polygon-scanning multiphoton microscopes have been demonstrated before, the FOV is relatively small (∼512 x 512 *μ*m) (***Li et al., 2018***, ***2017***). We used a polygon scanner to achieve a large scan angle at high scanning speed, which also reduces the complexity of the scanning engine when compared to the existing large FOV microscopes (***Sofroniew et al., 2016***; ***Stirman et al., 2016***; ***Ota et al., 2021***). The optical path and the footprint for the DEEPscope (∼3 ft x 3 ft x 1 ft, ***Figure 1—figure Supplement 1***) are nearly identical to a conventional multiphoton microscope. The polygon scanner has a larger aperture (9.5 mm) and more than twice the optical scan angle (∼60-degree peak-to-peak) than a resonant scanner. The polygon line rate (∼6 kHz) is > 6 times that of galvo scanners (< 1 kHz) at the same scan angle with ∼5 mm aperture size.

### Adaptive excitation and beamlet scanning for improving the multi-photon excitation efficiency

To reduce the average power required for simultaneous large FOV 2P and 3P imaging, an adaptive excitation scheme (***Li et al., 2020***) enabled by electro-optical modulators (EOMs) was used to block the laser in areas where large blood vessel shadows were located. ***Figure 1***d shows an image of cortical layer 6 (L6) with and without adaptive excitation. Fluorescence intensity increased during adaptive excitation because the effective power across the regions imaged was higher (224 mW versus 130 mW) even though the average power was lower (119 mW versus 130 mW).

To optimize excitation efficiency and improve imaging speed using a low repetition rate laser, we employed a beamlet scanning scheme with the PSF optimized to the size of neurons. A higher excitation efficiency improves the detection fidelity of calcium transients (d’) (***Wilt et al., 2013***). We underfilled the back aperture of the DEEPscope objective (∼85% filled). The axial resolution across the FOV was ∼5 *μ*m full-width half maximum (FWHM) (***Figure 1—figure Supplement 2***). This size of the PSF is close to the optimum for the size of neuron cell bodies, and approximately doubled d’ value by increasing the fluorescence signal ∼4 times when compared to a PSF with 2-*μ*m axial resolution (***Figure 1—figure Supplement 3***). This result is consistent with other studies (***Weisenburger et al., 2019***). In addition, we created a beamlet scanning scheme in which two pulses were scanned in two adjacent lines with a time delay of ∼20 ns (***Figure 1***c). Compared to scanning with a PSF with 10-*μ*m axial resolution, such a two-beamlet scanning scheme with two 5-*μ*m axial resolution PSF further increased d’ value by ∼10% (fluorescence signal by ∼20%) using the same amount of total pulse energy at the focal plane. Furthermore, the beamlets doubled the effective laser repetition rate and the line scanning speed, resulting in doubling both the spatial and the temporal resolution. More beamlets can be advantageous if higher pulse energy from the excitation source is available, providing greater gains in d’ value and higher spatial and temporal resolution (***Figure 1—figure Supplement 3***).

### Single-cell resolution, large-field-of-view 3P imaging in cortical L6 and deep imaging in CA1 through intact Cortex

We performed 3P large FOV structural imaging of neurons in a cortical column and the stratum pyramidale (SP) layer of the CA1 region of the hippocampus in a transgenic mouse (***Figure 2***, ***Figure 2***—***video 1*** and ***Figure 2***—***video 2***). We scanned a 3.23 mm x 3.23 mm field to cover most of the 3.5 mm diameter FOV of the microscope. We imaged a stack from 100 to 1,048 *μ*m in depth. GCaMP6s-expressing neurons in the SP layer appear at ∼872 *μ*m depth (***Figure 2***b). The external capsule (EC), where myelinated axons produce strong third harmonic generation (THG) signals, appeared to be curved across the large FOV and extended from ∼600 *μ*m to 1,048 *μ*m below the brain surface (***Figure 2***b). After the image acquisition, the digitally zoomed-in images and video (***Figure 2***b, ***Figure 2***c and ***Figure 2***—***video 1***) show the labeled neurons with clear nuclear exclusion in the whole cortical column and sub-cortical region. We also measured an axial resolution of ∼5 *μ*m in a vasculature-labeled mouse at a depth of 800 *μ*m below the brain surface (***Figure 2—figure Supplement 1***). ***Figure 2*** shows that 3P DEEPscope is capable of high-resolution, large FOV imaging deep within intact mouse brains.

**Figure 2.**
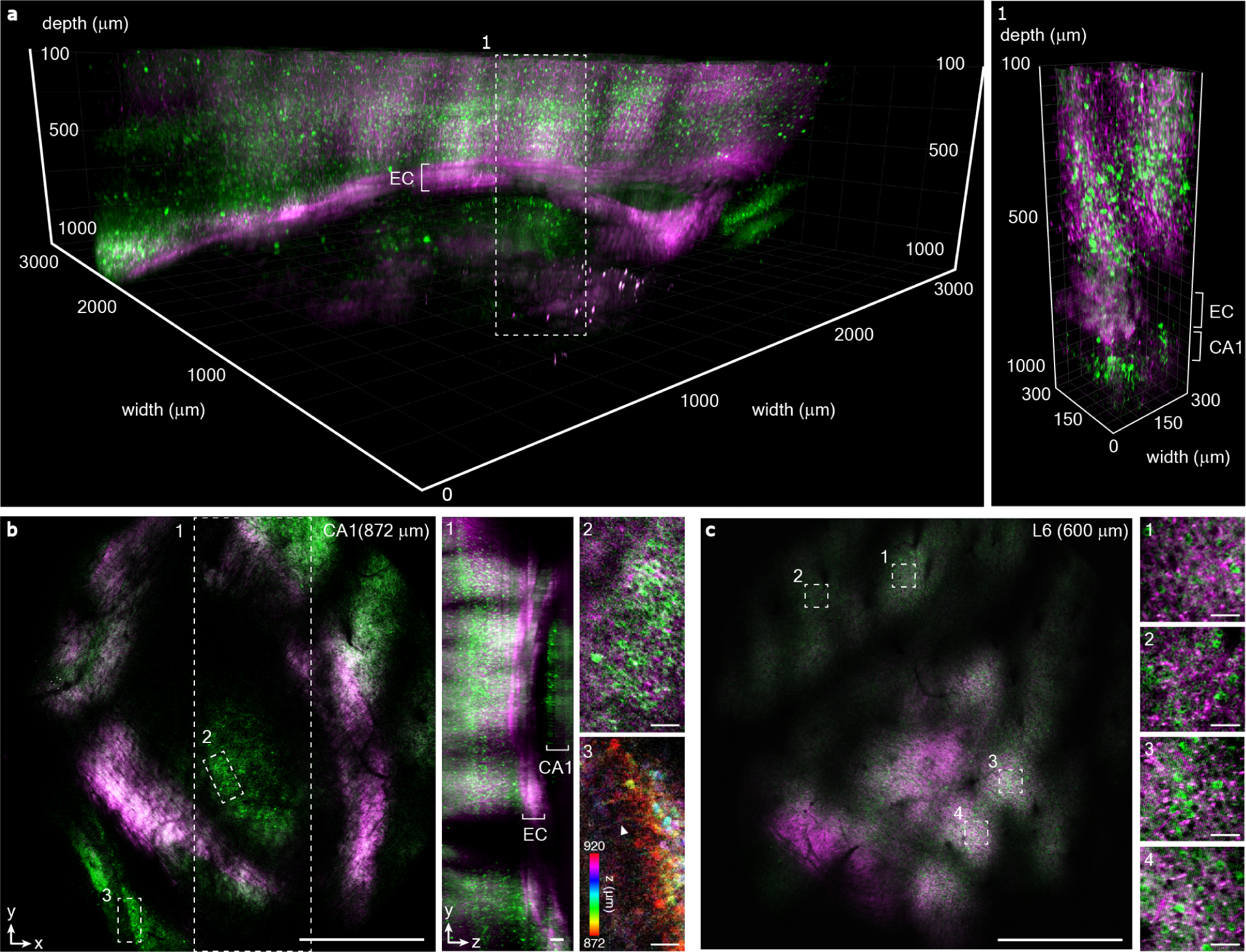
Deep and large FOV in-vivo imaging of mouse brain structures. (a) 3D rendering of GCaMP6s-expressing neurons by 3P DEEPscope from 100 to 1,048 *μ*m below the brain surface (CamKII-tTa/tetO-GCaMP6s mouse, male, 16 weeks old). The scanning field was ∼3.23 mm x 3.23 mm, acquired with 0.8-*μ*m pixel size and a z-step of 4 *μ*m. The integration time per imaging depth was 1.5 mins, 4 mins, and 10 mins at the depth of 100-900 *μ*m, 900-950 *μ*m, and 950-1,048 *μ*m respectively. Panel 1 shows the digitally zoomed-in volume of the 3D stack in a (indicated by the white dashed box) with 300 *μ*m FOV. (b) Selected 2D images from a of CA1 hippocampal neurons at an imaging depth of 872 *μ*m. Panel 1 shows the YZ maximum intensity projection of the dashed box 1 in b. Panels 2 and 3 show digitally zoomed-in images of the corresponding white dashed boxes in b. White arrowhead points to an axon. Scale bar in b, 1 mm. Scale bar in panel 1, 100 *μ*m. Scale bars in panels 2 and 3, 50 *μ*m. (c) Selected 2D image from a of layer 6 neurons at an imaging depth of 600 *μ*m. Panels 1-4 show digitally zoomed-in images of the corresponding dashed white boxes in c. Scale bar of c, 1 mm. Scale bars in panels 1 to 4, 50 *μ*m. EC: external capsule. **Figure 2—video 1.** Mouse brain imaging stack with 3.23 x 3.23 mm scan field from 100 to 1,048 *μ*m below the brain surface. The stack is acquired with 4116x4116 pixels per slice and 4 *μ*m step size. Image combines fluorescence signal (green) from neurons expressing GCaMP6s and THG signal (magenta) from a GCaMP6s transgenic mouse. **Figure 2—video 2.** 3D rendering of the imaging stack.

We imaged the spontaneous activity of GCaMP6s-expressing neurons in adult transgenic mice in the cortical L6 to demonstrate the capability of the 3P DEEPscope for large FOV activity imaging in the deep cortical regions. ***Figure 3***a shows the 3.23 mm x 3.23 mm scanned field and the digitally zoomed-in imaging sites in cortical L6 at 600 *μ*m depth, right above the EC. We used adaptive excitation to reduce the average excitation power from 224 mW to 119 mW while keeping the same pulse energy at the sample. We recorded neuronal activity traces from a large population of neurons (918 neurons) in an awake mouse (***Figure 3***b and ***Figure 3***—***video 1***) at a 4 Hz frame rate. ***Figure 3***c shows selected activity traces of the neurons. The raw photon counts of the activity recording are shown in ***Figure 3—figure Supplement 1***. We also imaged hippocampal SP neurons at 930 *μ*m depth with a FOV of 550 *μ*m x 550 *μ*m (***Figure 3***d and ***Figure 3***—***video 2***). The FOV for hippocampal imaging was reduced due to the limit on laser average power to prevent tissue heating. ***Figure 3***e shows selected activity traces of 42 neurons from the 503 hippocampal SP neurons.

**Figure 3.**
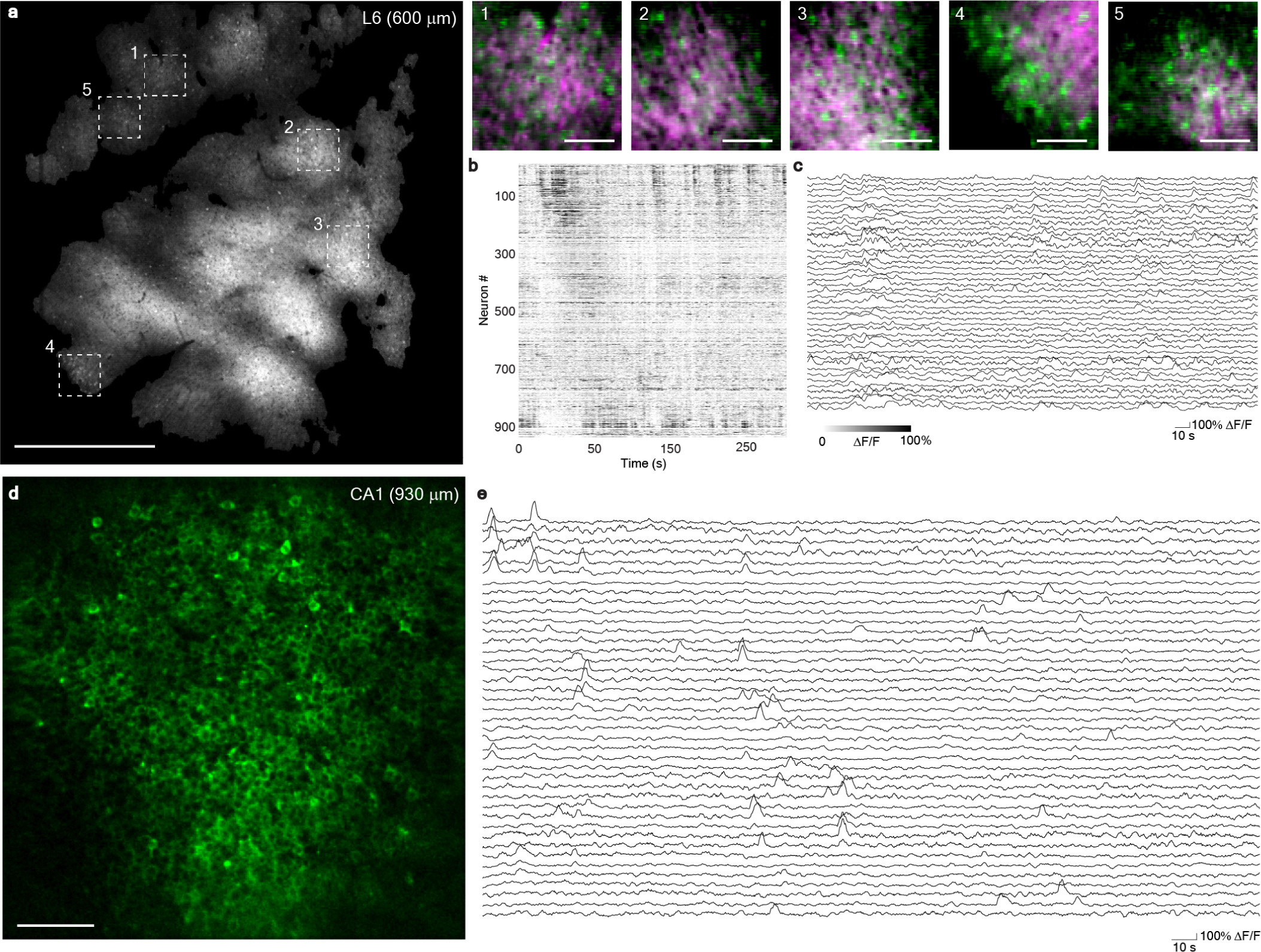
1320nm 3P activity imaging with DEEPscope (a) 3PM image of an activity recording site in an awake, GCaMP6s-expressing transgenic mouse (CamKII-tTa/tetO-GCaMP6s, male, 15 weeks old). The recording site is 600 *μ*m below the brain surface with a scanning field of ∼3.23 mm x 3.23 mm. Panels 1-5 show digitally zoomed-in images from the corresponding white dashed boxes in a. Scale bar, 1 mm. Scale bars in panels 1 to 5, 100 *μ*m. (b) Spontaneous activity traces recorded under awake conditions from 918 neurons in a. The average power on the brain surface was 119 mW. 53% of the scanned area was excited using the adaptive excitation scheme. The recordings were acquired with 1030x1030 pixels per frame (a pixel size of 3.14 *μ*m) and a frame rate of 4.0 Hz using two beamlets and the polygon scanner. (c) Selected activity traces from b. (d) Image of an activity recording site in an awake, GCaMP6s-expressing transgenic mouse (CamKII-tTa/tetO-GCaMP6s, female, 17 weeks old). The imaging took place in the sixth week after cranial window implantation. The recording site is at 930 *μ*m below the brain surface with a FOV of ∼*μ*m x 550 *μ*m. Scale bar, 100 *μ*m. (e) Selected spontaneous activity traces recorded from 42 neurons (out of 503 segmented neurons) in the FOV under awake condition. The repetition rate used for imaging was ∼1 MHz and the average power on the brain surface was 120 mW. The recordings are acquired with 512x512 pixels per frame, a pixel size of 1.06 *μ*m, and at 4.3 Hz frame rate using a galvo-galvo scanner. **Figure 3—video 1.** Spontaneous activity recorded from GCaMP6s-labeled neurons at cortical L6, 600 *μ*m below brain surface with 3.23 x 3.23 mm scan field at 4 Hz with 1030 x 1030 pixels per frame. Playback is sped up by 30x. Structural stack is shown to indicate the position of EC. Image combines fluorescence signal (green) from neurons expressing GCaMP6s and THG signal (magenta) from a GCaMP6s transgenic mouse. Activity recording is shown in gray scale. **Figure 3—video 2.** Spontaneous activity recorded from GCaMP6s-labeled neurons in SP layer of CA1 region of the hippocampus at 930 *μ*m below brain surface from a GCaMP6s transgenic mouse. Activity is recorded with a FOV 550 *μ*m x 550 *μ*m at 4.3 Hz frame rate. Playback is sped up by 30x. Kalman filtered video (left) and DeepCAD processed video (right) are shown.

### Dual excitation in 2P and 3P for large-field-of-view neuronal activity imaging at various depths

While 3P DEEPscope enables deep imaging in mice, 2P imaging achieves a higher volumetric imaging rate for shallow imaging depths due to its higher excitation repetition rate (***Wang and Xu, 2020***). Therefore, we designed the DEEPscope for excitation wavelengths of 910-930 nm, 1,030-1,050 nm, 1,240-1,380 nm, and 1,640-1,750 nm to enable both 2P shallow and 3P deep imaging. ***Figure 1—figure Supplement 4*** and ***Figure 1—figure Supplement 5*** show the transmission over the entire FOV for 920 nm and 1320 nm. ***Figure 4***a shows an imaging site of L5 neurons 480 *μ*m below the brain surface in an adult transgenic mouse. We recorded the neuronal activity traces of a large population of 2,000 neurons under awake conditions at a ∼6 Hz frame rate (***Figure 4***c and ***Figure 4***d). By reducing the FOV to ∼3.23 mm x 1 mm, we performed high spatial-resolution activity imaging (0.67 *μ*m pixel size) at a 4 Hz frame rate (***Figure 4***b). Nuclear exclusion and dendrites of the labeled neurons are clearly visible. We recorded the neuronal activity traces of 503 neurons (***Figure 4***e and ***Figure 4***f), which demonstrates that the DEEPscope can achieve high speed, large FOV, and high-resolution imaging in 2P imaging.

**Figure 4.**
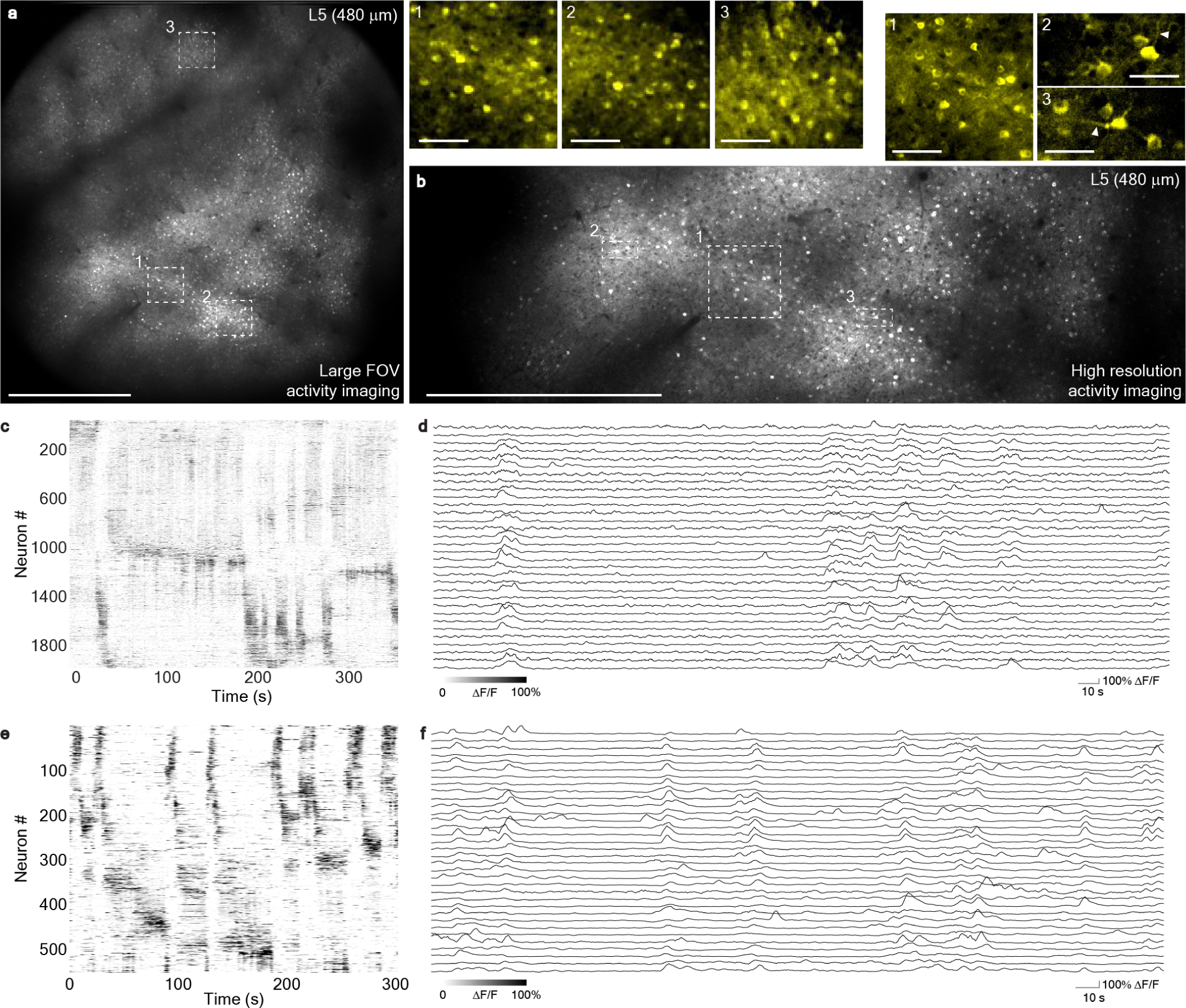
920nm 2P activity imaging with DEEPscope. Spontaneous activity recorded by 2P DEEPscope from cortical neurons (L5, 480 *μ*m below the surface of the brain) in a GCaMP6s-expressing transgenic mouse under awake condition (CamKII-tTA/tetO-GCaMP6s, male, 14-weeks old). The imaging took place in the second week after cranial window implantation. (a) Image of the recording site with a scanning field ∼3.23 x 3.23 mm acquired by 920 nm 2PM. Scale bar, 1 mm. Panels 1, 2 and 3 show digitally zoomed-in images of the corresponding white dashed boxes. Scale bar, 100 *μ*m. The average power on the brain surface was 130 mW with a pulse repetition rate of 80 MHz. The recordings were acquired with 1042x1042 pixels per frame, 3.14 *μ*m pixel-size and at 5.76 Hz frame rate. (b) High-resolution image of the recording site with a scanning field ∼3.23 x 1 mm of cortical L5 neuron acquired by 920 nm 2PM as a part of the imaging site in a. Scale bar, 1 mm. Panels 1, 2 and 3 show digitally zoomed-in images in the corresponding white dashed boxes in b. Panel 1 in b is at the same region as panel 1 in a. Scale bar in panel 1, 100 *μ*m. Scale bar in panel 2 and 3, 50 *μ*m. White arrowheads point to the dendrites. The average power on the brain surface was 130 mW with a repetition rate of 80 MHz. The recordings were acquired with 4862 x 1490 pixels per frame, 0.67 *μ*m pixel-size and at 4.0 Hz frame rate. (c) Spontaneous activity traces recorded under awake conditions from 2000 neurons in the FOV in a. (d) Selected activity traces from c. (e) Spontaneous activity traces recorded under awake conditions from 503 neurons in the FOV in b. (f) Selected activity traces from e.

We performed simultaneous six-plane 2P and 3P activity imaging of both shallow and deep cortical GCaMP6s-expressing neurons in adult transgenic mice. ***Figure 5*** shows 3.23 mm x 3.23 mm FOV 2P imaging of five focal planes at 320, 340, 360, 380, 400 *μ*m depth at a cycling rate of 2.2 Hz and 3P imaging at 600 *μ*m depth at a frame rate of 11 Hz. We were able to record neuronal activity traces from a large population of neurons (4,523 neurons) under awake conditions. ***Figure 5***b and ***Figure 5***c show spontaneous activity traces recorded by 920 nm 2PM under awake conditions from five focal planes in the FOV. ***Figure 5***d and ***Figure 5***e shows spontaneous activity traces recorded by 1320 nm 3PM.

**Figure 5.**
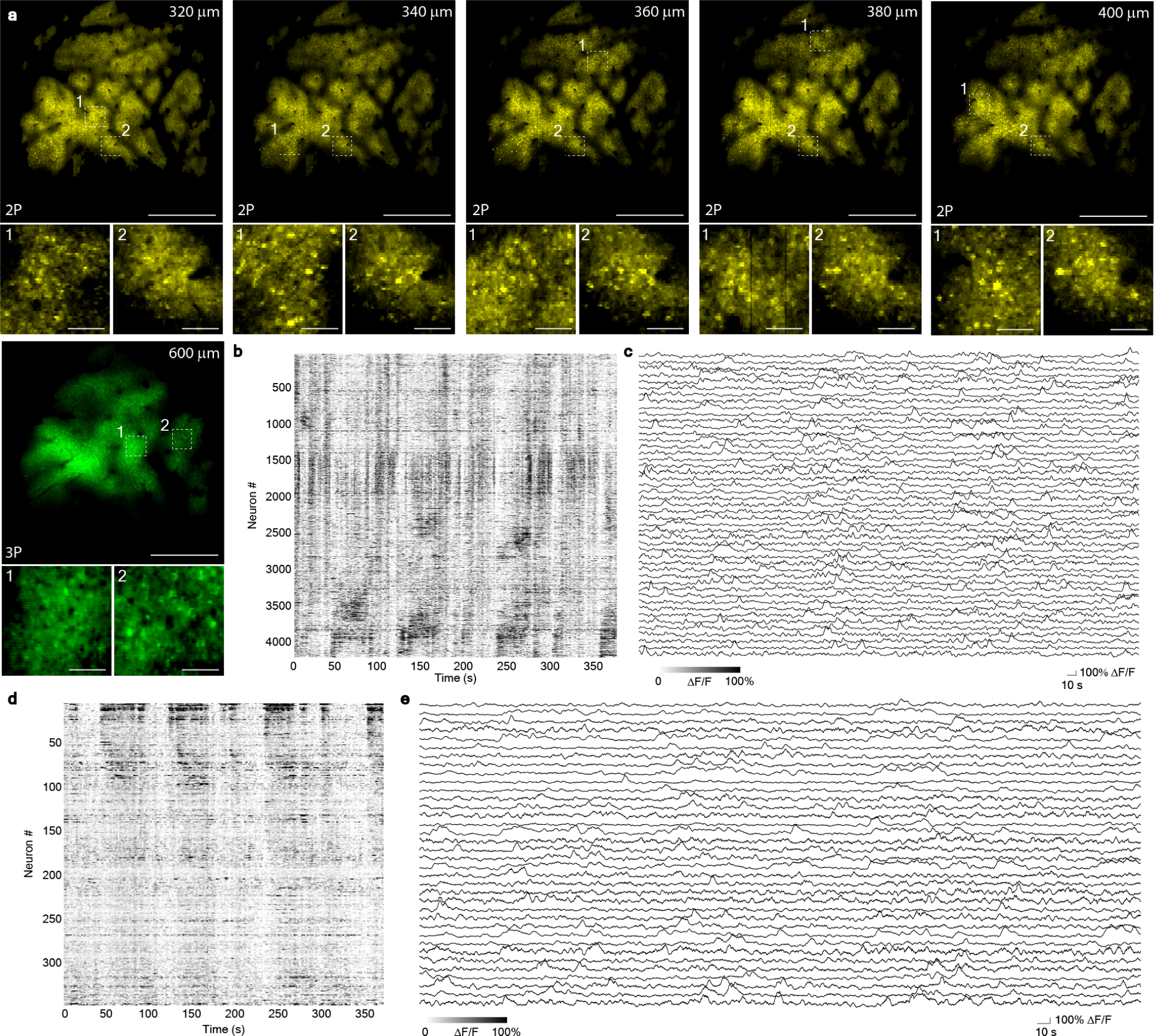
Large FOV neuronal activity recording in the shallow and deep cortex of GCaMP6s-expressing transgenic mouse. (a) Images of six focal planes acquired simultaneously by 1320 nm 3PM (green) and 920 nm 2PM (yellow) in an awake, GCaMP6s-expressing transgenic mouse (CamKII-tTa/tetO-GCaMP6s, female, 17-weeks old). The activity recording sites were 320, 340, 360, 380, 400 and 600 *μ*m below the brain surface with a scanning field of ∼3.23 x 3.23 mm. Scale bar, 1 mm. Panels 1 and 2 show digitally zoomed-in images from the corresponding white dashed boxes. Scale bar, 100 *μ*m. (b) Spontaneous activity traces recorded by 920 nm 2PM under awake conditions from 4183 neurons (total from the five focal planes) in the FOV in a. (c) Selected activity traces from b. (d) Spontaneous activity traces recorded by 1320 nm 3PM under awake conditions from 340 neurons in the FOV in a. (e) Selected activity traces from d. For 3P imaging, the average power on the brain surface was 97 mW. 46% of the scanned area was excited using the adaptive excitation scheme. The recordings were acquired with 1030x1030 pixels per frame, a pixel size of 3.14 *μ*m and a frame rate of 11 Hz using two beamlets. For 2P imaging, the laser repetition rate used for imaging was 80 MHz, and the average power on the brain surface was 75 mW. 62% of the scanned area was excited using the adaptive excitation scheme. The recordings were acquired with 1030 x 515 pixels per frame, a pixel size of 3.14 x 6.28 *μ*m and a volume rate (i.e., the cycling rate among the five focal planes) of 2.2 Hz.

### 3P large FOV imaging of adult Zebrafish brain

The DEEPscope can be applied to other animal models as well. We performed 3P large FOV imaging of an adult zebrafish brain in-vivo (***Chow et al., 2020***). We imaged a stack from 0 to 1,090 *μ*m in depth (***Figure 6***, ***Figure 6***—***video 1*** and ***Figure 6***—***video 2***). Individual GCaMP6s-expressing nuclei were clearly visible in the telencephalic, optic tectum, and cerebellar regions and the THG signal revealed bone structures and fiber tracts. The entire olfactory bulbs, the olfactory nerves, and part of the olfactory epithelium were also visible.

**Figure 6.**
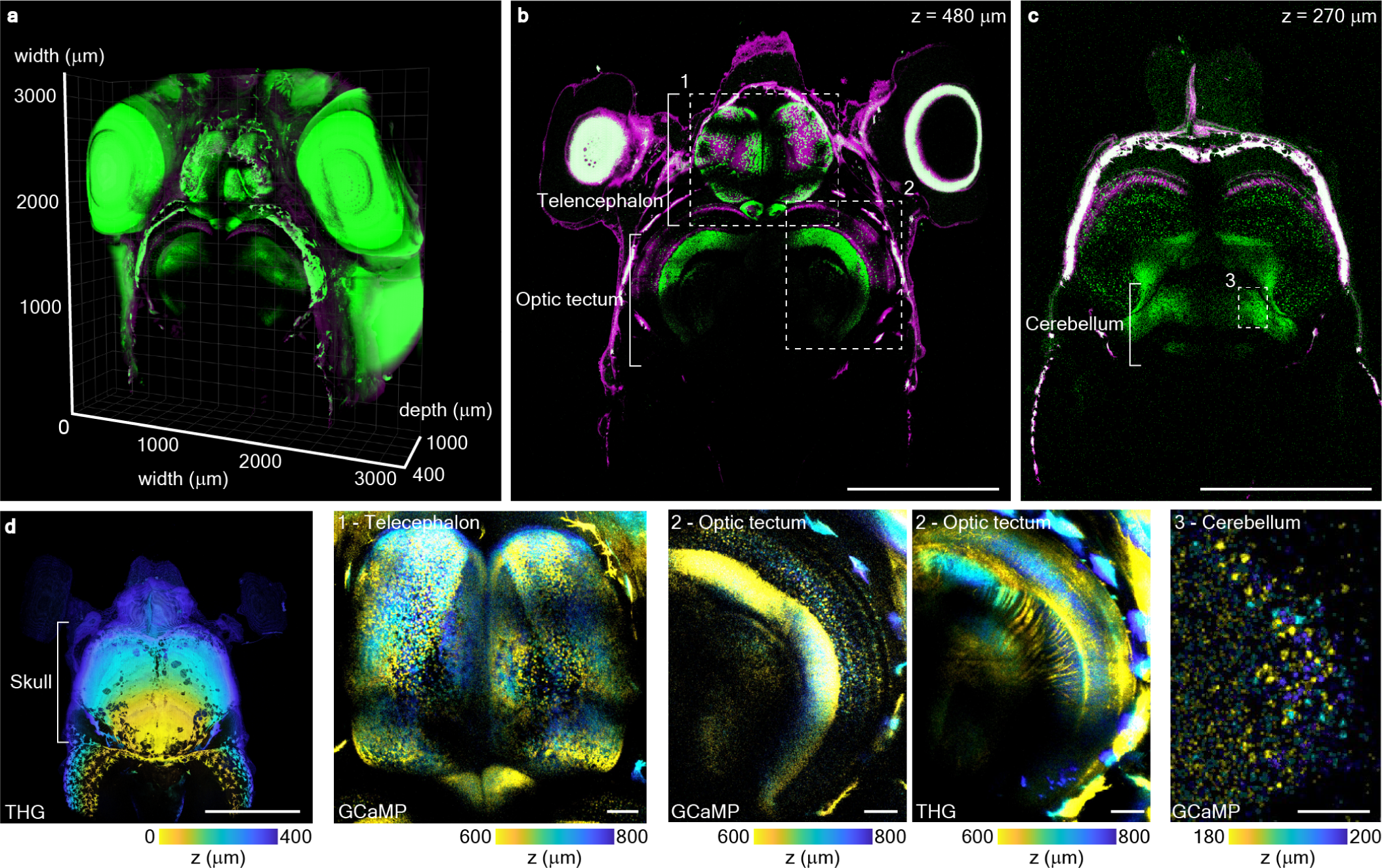
Brain-wide imaging of a living, intact adult zebrafish. (a) 3D rendering of structural imaging of GCaMP6s-expressing nuclei (green) and THG signal (magenta) from regions with myelinated fibers obtained by 3P DEEPscope from 400 *μ*m to 1090 *μ*m below the surface of the skull bone of an adult zebrafish (Tg(elavl3::H2B-GCaMP6s, male, 9 months old, 15.4 mm standard lengths). The scanning field is ∼3.23 x 3.23 mm, acquired with 0.8-*μ*m pixel size and 10-*μ*m z-step using the polygon scanner. (b) Image of the zebrafish at an imaging depth of 480 *μ*m, showing the brain region telencephalon and optic tectum. GCaMP6s-expressing nuclei are shown in green and THG signal is shown in magenta. Scale bar, 1 mm (c) Image of the zebrafish at an imaging depth of 270 *μ*m, showing the brain region cerebellum. GCaMP6s-expressing nuclei are shown in green and THG signal is shown in magenta. Scale bar, 1 mm (d) THG image of the zebrafish at the imaging depths of 0 - 400 *μ*m, showing the skull bone. The color map shows the imaging depth. Scale bar, 1 mm. Panels 1, 2 and 3 show digitally zoomed-in images of the corresponding dashed white boxes in b and c with a color map showing the imaging depth. Scale bar, 100 *μ*m. The integration time per imaging depth was 0.5 min, 1 min and 1.7 mins at the depths of 0-750 *μ*m, 760-900 *μ*m and 910-1090 *μ*m, respectively. **Figure 6—video 1.** Zebrafish imaging stack with 3.23 x 3.23 mm scan field from 0 to 1,090 *μ*m below the top surface of the skull bone. The stack is acquired with 4123x4123 pixels per slice and 10 *μ*m step size. Image combines fluorescence signal (green) from neurons expressing GCaMP6s and THG signal (magenta) from neural fibers. **Figure 6—video 2.** 3D rendering of the imaging stack from depths of 400 *μ*m to 1,090 *μ*m.

## Discussion

The Imaging depth, FOV, and speed are ultimately limited by the achievable fluorescence signal. The use of adaptive excitation, which involves simply blocking the lasers with EOMs in the uninformative parts of the sample suggests potential for further increasing the excitation efficiency. A customized adaptive excitation source (AES), for example, illuminates the neurons with a high instantaneous repetition rate. It can increase the fluorescence signal by more than 10-fold without increasing the average power as shown in a previous paper (***Li et al., 2020***). New transgenic mice with brighter jGCaMP7s or jGCaMP8s indicators will further increase fluorescence signal by > 4-fold (***Bounds et al., 2021***; ***Dana et al., 2019***; ***Zhang et al., 2023***) for single and multiple action potentials. The combination of the DEEPscope, the AES, and brighter calcium indicators will further improve the capability for deep, fast, and large FOV imaging of neuronal activity.

Using commercially available 3P and 2P laser systems, a polygon scanner, and EOMs, we have demonstrated a number of methods to significantly improve the field-of-view and imaging speed for both 2P and 3P imaging systems. By employing adaptive excitation, we can increase the number of recorded neurons without increasing the average power. By generating spatially dispersed beamlets, we can also enhance the scanning rate beyond the limits of the scanners while preserving high spatial resolution, strong multiphoton signal, and a large scanning FOV. The combination of high imaging speed and large volume offered by the DEEPscope will allow simultaneous monitoring of the dynamics of a large population of cells across a large FOV in different kinds of tissues.

## Methods and Materials

### Dual Excitation with adaptive Excitation Polygon-scanning multiphoton microscope (DEEPscope)

#### Excitation source and adaptive excitation

The excitation source for 3PM was a noncollinear optical parametric amplifier (NOPA, Spectra-Physics) pumped by an regenerative amplifier (Spirit, Spectra-Physics). The NOPA operated at a center wavelength of 1320 nm and provided an average power of ∼2 W at 2 MHz repetition rate (∼1 *μ*J per pulse). It was paired with an electro-optic modulator (EOM) (360-40-03, Conoptics) for adaptive excitation (***Li et al., 2020***). A two-prism (SF11 glass) compressor was used to compensate for the normal dispersion of the optics in the light source and the microscope. The pulse duration (measured by second-order interferometric autocorrelation) under the objective was ∼60 fs after dispersion compensation.

The excitation source for 2PM was a mode-locked Ti:Sapphire laser (Chameleon, Coherent). The Ti:Sapphire laser operated at a center wavelength of 920 nm and provided an average power of ∼1.6 W at 80 MHz repetition rate. It was paired with an EOM (350-160, Conoptics) for adaptive excitation. A two-prism (SF11 glass) compressor was used to compensate for the normal dispersion of the optics in the light source and the microscope. The pulse duration (measured by second-order interferometric autocorrelation) under the objective was ∼90 fs after dispersion compensation.

#### Beamlet generation delay line

A beamlet generation delay line (***Figure 1***) was used to split a laser pulse at 1320 nm into two pulses (beamlets) with ∼20 ns time delay, increasing the effective laser repetition rate from 2 MHz to 4 MHz. The delay line was arranged as a loop with four one-inch dielectric mirrors (102077, Layertec) and a polarizing beam splitter cube (PBS) (PBS103, Thorlabs). The PBS served as the input and output port of the delay line. The power distribution of each beamlet was controlled by adjusting the half-wave plate (HWP) (WPH05M-1310, Thorlabs) before the PBS. Inside the loop, two mirrors were tilted to adjust the angular separation between the beamlets. The angularly separated beamlets converge on the polygon scanner in a plane that was perpendicular to the scanning direction. An 8-f system was placed inside the loop to compensate for the difference in beam divergence between the beamlets. Two co-planar foci that were ∼3 *μ*m apart were created along the slow axis (***Figure 1—figure Supplement 9***). The fluorescence signal generated by the two beamlets is demultiplexed temporally and forms the pixels in the corresponding lines in the image. The crosstalk of the fluorescence signals between the two beamlets was measured to be ∼5%, which was caused by the data acquisition bandwidth of the system (***Figure 1—figure Supplement 6***).

#### DEEPscope setup

We developed a multiphoton microscope that enables 3.5 mm diameter large FOV imaging (***Figure 1—figure Supplement 7***) using a custom-designed scan lens (f = 60 mm), tube lens (f = 200 mm), and objective lens (f = 15 mm) (Special Optics, Navitar). The objective lens is water immersion and has a maximum excitation numerical aperture (NA) of 0.75 and a collection NA of 1.0 (***Figure 1—figure Supplement 8***) with a 2.8 mm working distance. The objective was underfilled with 1/e^2^ beam diameter of ∼17 mm to achieve an axial resolution (FWHM) of ∼5 *μ*m and lateral resolution (FWHM) of ∼0.7 *μ*m across the FOV (***Figure 1—figure Supplement 2*** and ***Figure 1—figure Supplement 9***) for both 2P excitation at 920 nm and 3P excitation at 1320 nm. A 9-mm aperture polygon scanner and a 10-mm aperture galvo-mirror (Saturn 9B, ScannerMax) were conjugated using two custom-designed scan lenses (f = 60 mm) (Special Optics, Navitar) (***Figure 1***). The customized polygon scanner (PT60SRG, Nidec Copal Electronics) has 12 facets, each with 17.5 mm x 9.5 mm clear aperture. Since beam truncation occurs at the edge of each facet, the fill fraction of the polygon scan was 70% both temporally and spatially to ensure uniform transmission across the field of view. A 42-degree peak-to-peak optical scan angle was used to achieve a 3.23 mm linear scan field. The polygon scanner we used is capable of scanning a 60-degree peak-to-peak optical angle with a variable rotation speed of 7,000 – 30,000 revolutions per minute (RPM), i.e., 1.4 kHz – 6 kHz line rate. Polygon scanner with even higher rotation speed, e.g., 55,000 RPM, is possible from different manufacturers, potentially further increasing the scan speed to an 11 kHz line rate.

For hippocampus activity imaging (***Figure 3***d), a 5 mm-aperture non-conjugated galvo-galvo scanner (Saturn 9B, ScannerMax) was used since the FOV was small. This configuration shared the same lenses and effective excitation NA (***Figure 1—figure Supplement 2***) as the polygon-galvo configuration. The galvo-galvo scanner could achieve ∼1 kHz line rate at ∼40-degree peak-to-peak optical scan angle.

A fast remote focusing module (***Figure 1***) enabled tunning of the 2P focal plane away from the 3P focal plane. In the 2P beam path, an HWP (WPH05M-915, Thorlabs) and PBS (PBS255, Thorlabs) directed the beam onto a quarter wave plate (QWP) and the remote focusing objective (ROBJ) (LMH-50X-850, Thorlabs). The QWP and ROBJ were double passed after reflection from a small mirror (PF03-03-P01) and a custom-made adapter (∼20g total weight), which were mounted on a voice coil (LFA 2010; Equipment Solutions). The back aperture of the ROBJ was conjugated to the back aperture of the imaging objective. The axial resolution at different focal plane positions during remote focusing is shown in ***Figure 1—figure Supplement 11***.

For signal collection, the fluorescence and third harmonic generation (THG) signal were epi-collected through the imaging objective lens and immediately reflected by a 77 mm x 108 mm dichroic beam splitter (FF705-Di01, Semrock) to a detection system (Special Optics, Navitar) that was custom-designed to achieve a high collection efficiency. Another 77 mm x 108 mm dichroic mirror (FF470-Di01, Semrock) split the signal into two channels: one for the fluorescence signal emitted from GCaMP6s and the other for the THG signal. One-inch optical filters 520/70 (FF01-520/70, Semrock) and 435/40 (FF02-435/40, Semrock) were used for the fluorescence and THG channels, respectively. The signals were detected with two 14x14 *mm*^2^ effective sensor area GaAsP photomultiplier tube (H15460-40) that was customized for low dark-count and with the built-in preamplification unit removed. The detection efficiency of the system was estimated to be ∼3.5% for fluorescence imaging across the 3.5 mm FOV at an imaging depth of 900 *μ*m (using an established empirical model (***Beaurepaire and Mertz, 2002***) and the measurement result in ***Figure 1—figure Supplement 8***).

The PMT current was converted to voltage by a transimpedance amplifier (HCA-200M-20K-C, Femto). Analog-to-digital conversion was done by a data acquisition card at a sampling clock of 123 MHz (vDAQ, Vidrio), triggered by the Spirit-NOPA system at 1.96 MHz with a 63x electronic multiplier. Light shielding was carefully done to achieve a dark count of ∼1000 photons per second under the usual imaging environment. The acquisition system achieved shot-noise-limited performance. 3P signals for the two beamlets were acquired with two acquisition gates for time demultiplexing. A customized ScanImage 2021 (Vidrio) running on Matlab (MathWorks) was used to place the 3P signals from each beamlet into two virtual channels. These channels were interleaved with a custom Matlab script after image acquisition. 2P signal was placed into a separate virtual channel. A translation stage was used to move the sample (M-285, Shutter Instrument). For depth measurement, the slightly larger index of refraction in brain tissue relative to water resulted in a slight underestimate (5–10%) of the actual imaging depth within the tissue, because the imaging depths reported here are the raw axial movement of the sample (***Wang et al., 2018***).

#### Two-photon and three-photon temporal multiplexing

Simultaneous 2P and 3P excitation were achieved by temporal multiplexing of the 920 nm Ti:Sapphire laser and the 1320 nm Spirit-NOPA. The setup is similar to the one described in a previous study (***Ouzounov et al., 2017***). Briefly, the two excitation beams were combined with a 980 nm long pass dichroic mirror (DMLP1000R, Thorlabs) and passed through the same scanners. They were spatially separated into different focal planes by using the remote focusing module. The 920 nm laser was intensity modulated with an EOM, which was controlled by a transistor-transistor logic (TTL) gate signal. The TTL signal was generated from a signal generator (SDG2042X, Siglent) that was triggered by the Spirit-NOPA laser. The EOM had high transmission for 360 ns between two adjacent Spirit-NOPA laser pulses (or pulse pairs for the two beamlets) that were ∼500 ns apart.

#### Adaptive excitation

A structural image was first obtained from conventional raster scanning. Then Gaussian filters and median filters were used to remove sharp features in the image. The regions for imaging were selected by using the top 80% of pixel intensity in the structural image, which effectively excludes regions of the blood vessel shadows. The positions of the areas for imaging were then converted into a digital time sequence for each scan line and sent to an arbitrary waveform generator (AWG). The AWG, triggered by the line trigger from ScanImage, controlled a Pockels cell to transmit laser power only to the selected areas within the FOV. Adaptive excitation for 3PE was achieved with one AWG (PXI-5412, National Instrument). Adaptive excitation for 2P/3P multiplexing was achieved with two AWGs (PXI-5421 and PXI-5412, National Instrument) to accommodate different selected areas at different imaging depths. The AWGs were configured to have a sampling rate of 5 MHz. For 3PM, the time sequence from the AWG was directly sent to the Pockels cell driver (302A, Conoptics). For 2PM, the time sequence from the AWG was combined with the TTL gate signal (see the section on Two-photon and three-photon temporal multiplexing) with an AND gate (TI SN74HC08N, Texas Instruments) before sending it to the Pockels cell driver (25D, Conoptics) (***Figure 1—figure Supplement 10***).

#### Image acquisition parameters

All acquisition parameters for structural and functional imaging are summarized in Table 1.

**Table 1.** Acquisition parameters for deep and large FOV two- and three-photon imaging shown in this work.

| Figure | Imaging modality | Imaging depth ($\mu\text{m}$) | Imaging FOV ($\text{mm}^2$) | Power on sample (mW) | Duty cycle of adaptive excitation (%) | Number of pixels | Laser repetition rate (MHz) | Number of beamlets | Frame Rate (Hz) |
| --- | --- | --- | --- | --- | --- | --- | --- | --- | --- |
| 2 | 3P | 100–1048 | 3.23x3.23 | <130 | 53 | 4116x4116 | 2 | 2 |  |
| 3a | 3P | 600 | 3.23x3.23 | 119 | 53 | 1030x1030 | 2 | 2 | 4 |
| 3d | 3P | 930 | 5.50x5.50 | 120 | 53 | 512x512 | 1 |  | 4.3 |
| 4a | 2P | 480 | 3.23x3.23 | 130 | 53 | 1042x1042 | 80 |  | 5.7 |
| 4b | 2P | 480 | 3.23x3.23 | 130 | 53 | 4862x1490 | 80 |  | 4 |
| 5 | 3P | <600 | 3.23x3.23 | 97 | 46 | 1030x1030 | 2 | 2 | 11 |
| 6 | 3P | 380,400 | 3.23x3.23 | <75 | 62 | 4123x4123 | 2 |  | 2.2 |
| Videos |  |  |  |  |  |  |  |  |  |
| Figure 2—video 1, Figure 2—video 2 |  |  | Same parameters as Figure 2 |  |  |  |  |  |  |
| Figure 3—video 1 |  |  | Same parameters as Figure 3a |  |  |  |  |  |  |
| Figure 3—video 2 |  |  | Same parameters as Figure 3d |  |  |  |  |  |  |
| Figure 6—video 1, Figure 6—video 2 |  |  | Same parameters as Figure 6 |  |  |  |  |  |  |

### Image processing for structural recording

Structural imaging was normalized by the linear transform of pixel intensities to saturate the brightest 0.1-0.5% pixels in each frame. Three-dimensional reconstruction of the stacks was rendered in Imaris.

### Image processing and data analysis for activity recording

Motion corrections were performed using Suite2p (***Pachitariu et al., 2016***). Neuron segmentation was done using Suite2p or manually with ImageJ. Extractions of fluorescence time traces (F) were done with Suite2p or a custom Matlab script. In Matlab, traces (F) were filtered with a moving average of window size of 2s. Baselines of the traces (F_0_) were determined by excluding the spikes and their rising and falling edges. Traces (F) were normalized according to the formula (F-F_0_)/F_0_. For the visual representation of calcium activities in ***Figure 3***—***video 1***, the raw image sequence was processed by Kalman filter with a gain of 0.7. For ***Figure 3***—***video 2***, the raw image sequence was processed by moving average of 4 frames and Kalman filter with a gain of 0.8 (left) and by DeepCAD(30) (right).

### Excitation efficiency optimization using discriminability index (d’)

The discriminability index (d’) for calcium transient detection using the beamlet scanning schemes (***Figure 1—figure Supplement 3***) was calculated according to equations for three-photon excited fluorescence in a Gaussian focus (***Wang et al., 2020***; ***Xu and Webb, 2002***). All calculations were performed assuming a spherical region of interest (ROI) with a radius of 5 *μ*m. Excitation inside the ROI yields a fluorescence signal while excitation outside the ROI yields background. To maximize power efficiency, we assumed 100% labeling density to penalize excitation outside the ROI and to reduce neuropil contamination. The refractive index of the medium was 1.33. The center wave-length was 1320 nm and the laser pulse width was 60 fs (FWHM). Three-photon cross-section of GCaMP6s (3 x 10^−82^ cm^6^s^2^) was used to calculate the fluorescence signal and the saturation pulse energy.

### Animal surgery and in-vivo calcium imaging of awake mice

All animal experiments and housing procedures were conducted in accordance with Cornell University Institutional Animal Care and Use Committee guidance. Chronic craniotomy was performed on mice according to the procedures described in the previous work (***Ouzounov et al., 2017***). Briefly, a window of 5 mm diameter was created, centering at ∼2.5 mm lateral and ∼2 mm caudal from the bregma point over the somatosensory cortex. Calcium imaging was performed on transgenic animals with GCaMP6s-expressing neurons (male and female, 15-18 weeks, CamKII-tTA/tetO-GCaMP6s). The spontaneous calcium activity imaging was performed on awake animals. The imaging took place 2-8 weeks after cranial window implantation.

### Vasculature imaging for resolution measurement

The mice were anesthetized with isoflurane (1–1.5% in oxygen, with breathing frequency maintained at 1 Hz) and placed on a heat blanket to maintain body temperature at 37.5°C. Eye ointment was applied. The vasculature of the transgenic mouse (TIT2L-GC6s-ICL-tTA2, female, 25 weeks) was labeled via retro-orbital injection of fluorescein (25mg of dextran conjugate dissolved in 200 mL of sterile saline, 70-kDa molecular weight; D1823, Invitrogen).

### Animal surgery and in vivo calcium imaging of zebrafish

Adult zebrafish (*Danio rerio*) (Tg(elavl3::H2B-GCaMP6s)), male, 9 months postfertilization (***Vladimirov et al., 2014***) were used. Small to medium-sized fish were chosen (standard lengths (tip of the head to the base of tail) 15.4 mm) for whole brain imaging. The fish is prepared similar as described in (***Chow et al., 2020***). To be brief, the fish were anesthetized with 0.2 mg mL^−1^ tricaine solution (pH 7.2). Then, 2 mL pancuronium bromide (0.4 mg mL-1 in Hanks) was retro-orbitally injected to paralyze the fish. The fish was then transferred to a petri dish with a ‘V’-shaped mounting putty (Loctite) to support the fish with the dorsal side up. A drop of the anesthetic bupivacaine was placed on the head. A small strip of putty was gently placed over the back of the fish to secure it. Fish were perfused through the mouth with an ESI MP2 Peristaltic Pump (Elemental Scientific) at a rate of 2 mL min ^−1^ with oxygenated fish system water during the experiment.

## Acknowledgments

We thank members of the Xu research group, especially Bo Li, Xusan Yang, and Kibaek Choe for their help and valuable discussions. We thank Spencer Smith and Che-Hang Yu for the discussion of optics design. We thank Katherine Strednak, Emily Silvela, and Faith Burgus from Cornell Center for Animal Resources and Education for their animal care service. We thank Karl Termini from the Cornell glass shop for his help in glass grinding. We thank Robert Page, Stanley McFall, Chris Cowulich, and Jeffrey Koski from the Cornell machine shop for their help in machining. We thank the Cornell Institute of Biotechnology for providing a workstation for 3D rendering. We thank Vidrio, Special Optics, Nidec Copal Electronics, Hamamatsu Photonics, ScannerMax, Eksma Optics, and AVR optics for customized parts.

## Author contributions

Conceptualization: AM, TW, CX

Methodology: AM, TW, SZ, DGO, CS, CW

Investigation: AM, KEK, JF, DW

Visualization: AM

Supervision: CX

Writing—original draft: AM

Writing—review and editing: TW, CX

---

**Figure 1—figure supplement 1.**
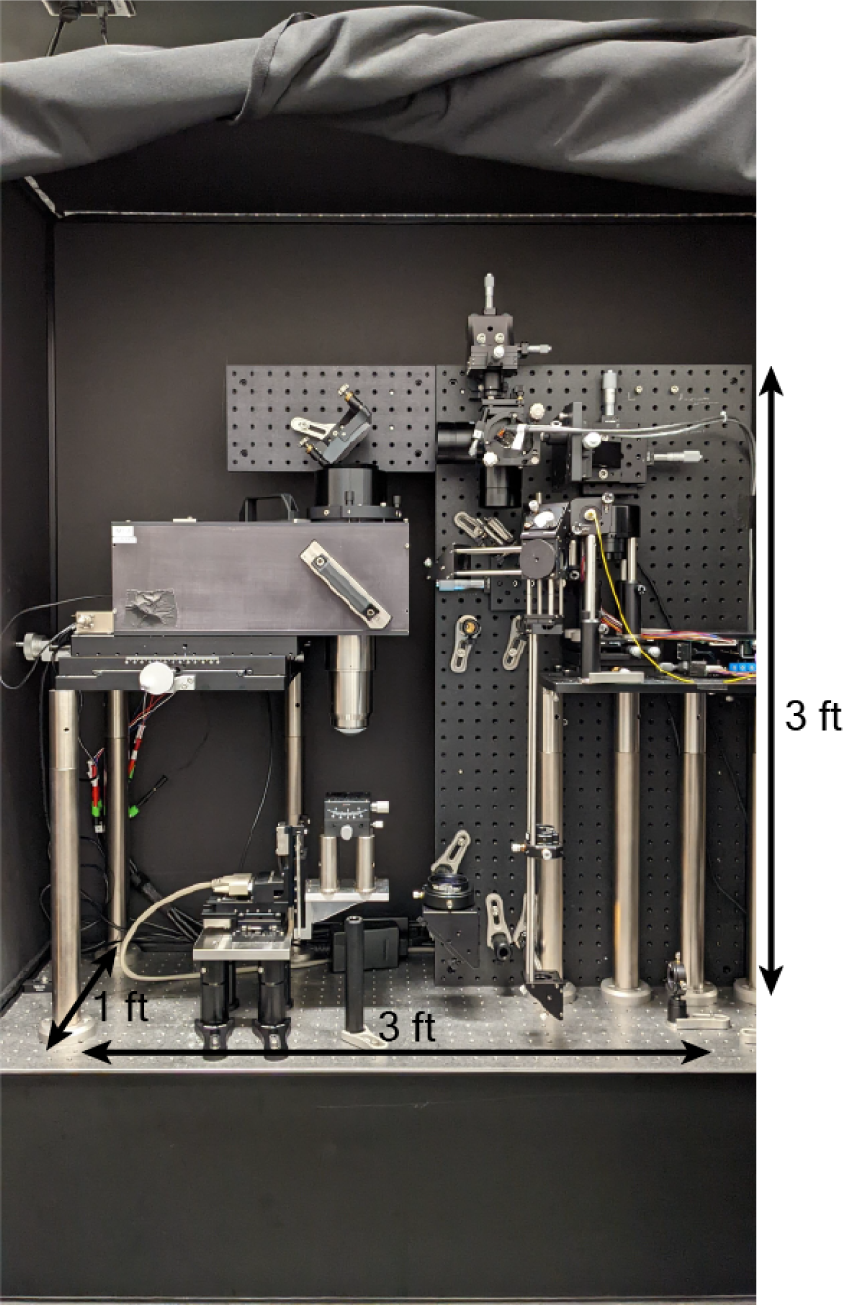
Photo of DEEPscope. The dimensions of the DEEPscope are ∼3 ft x 3 ft x 1 ft feet.

**Figure 1—figure supplement 2.**
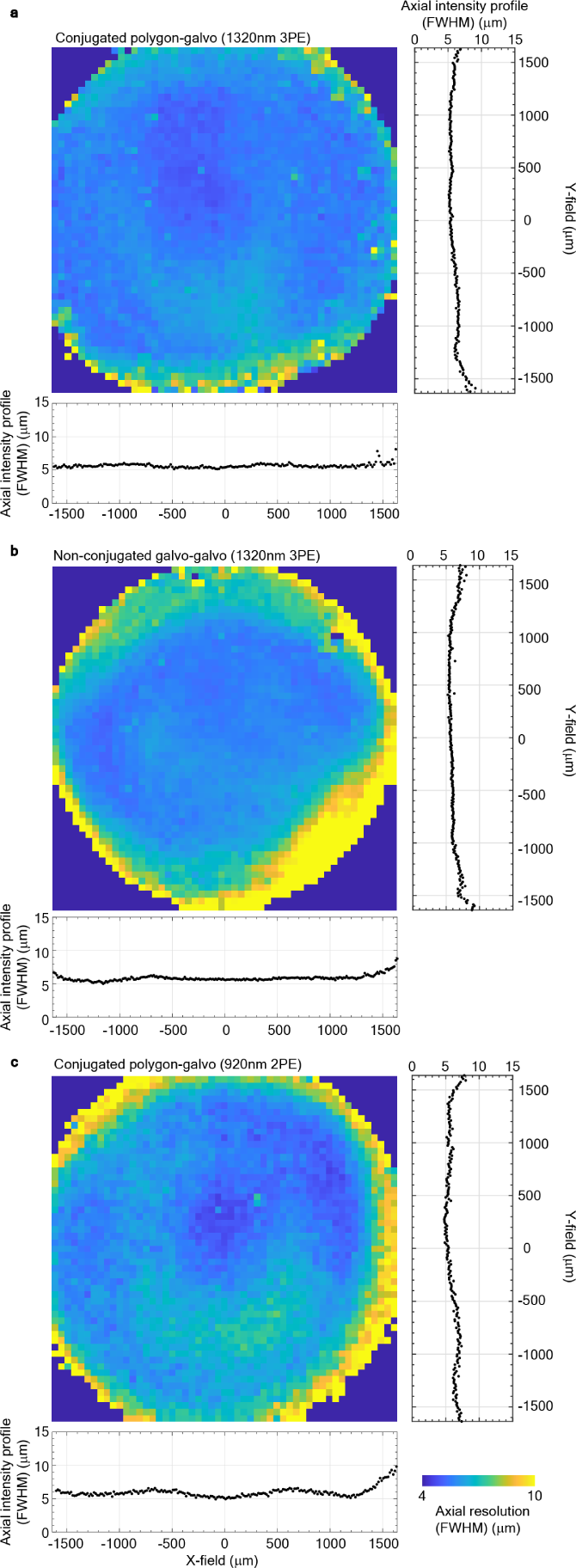
Characterization of the axial resolution of DEEPscope across the field-of-view by axial scanning of a 500-nm thin fluorescent film of Rhodamine B (RhB) dye.**(a)** FWHM of the axial intensity profiles of DEEPscope across the field-of-view measured by axial scanning of the thin fluorescent film with 1320 nm three-photon excitation (3PE). The scan engine consists of a pair of conjugated polygon-scanner and a galvo-scanner. **(b)** FWHM of the axial intensity profiles of DEEPscope across the field-of-view measured by axial scanning of the thin fluorescent film with 1320 nm three-photon excitation (3PE). The scan engine consists of a pair of closely spaced galvo-scanners without relay lenses between the two galvo-scanners.**(c)** FWHM of the axial intensity profiles of DEEPscope across the field-of-view measured by axial scanning of the thin fluorescent film with 920 nm two-photon excitation (2PE). The scan engine consists of a pair of conjugated polygon-scanner and galvo-scanner.

**Figure 1—figure supplement 3.**
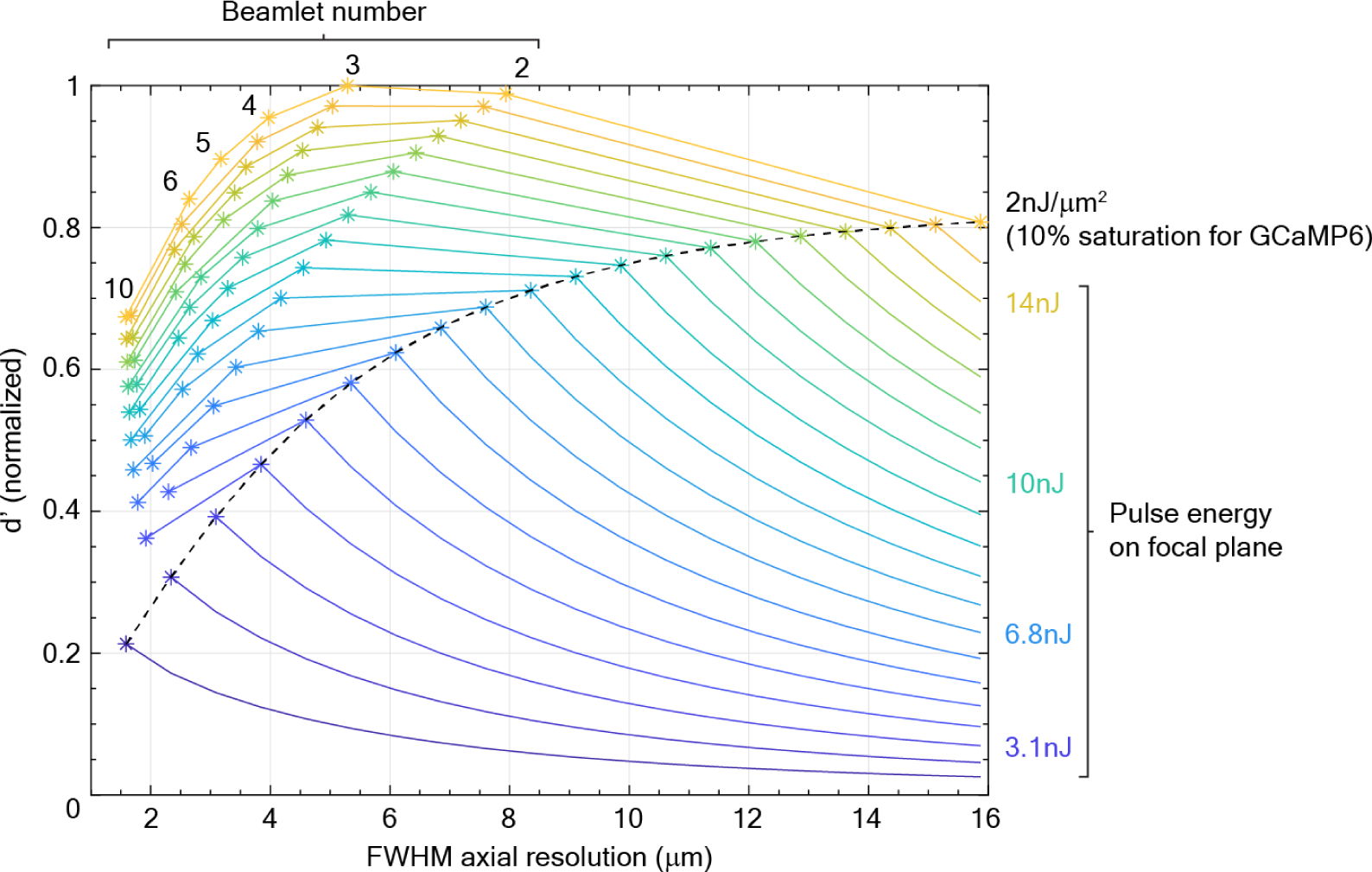
Simulation result of the relative d’ value with different excitation parameters for 1320 nm 3P imaging. Limited by the excitation source and the loss of the optical system, to image at 600 *μ*m depth, the best performance is realized by a two-beamlets approach with total pulse energy of ∼10 nJ at the focal plane. Normalized d’ values are shown for the excitation of a 5-*μ*m radius sphere using different focal spot size (i.e., FWHM axial resolution) and pulse energy at the focal plane. Pulse energies are fixed for each line with the same color. Black dashed line shows the target pulse fluence at 2 nJ/*μ*m^2^ with a 60 fs pulse at the focal point, which corresponds to 10% excitation saturation of GCaMP6 (i.e., the probability of excitation per pulse per molecule is 10% at the focal point). Each asterisk shows the d’ value when splitting the beam into different number of beamlets. The color of the asterisks corresponds to the sum of the pulse energy of all the beamlets (indicated on the right side of the graph). For all asterisks, the pulse fluence of each beamlet is fixed at 2 nJ/*μ*m^2^ (i.e., the target pulse fluence set by excitation saturation).

**Figure 1—figure supplement 4.**
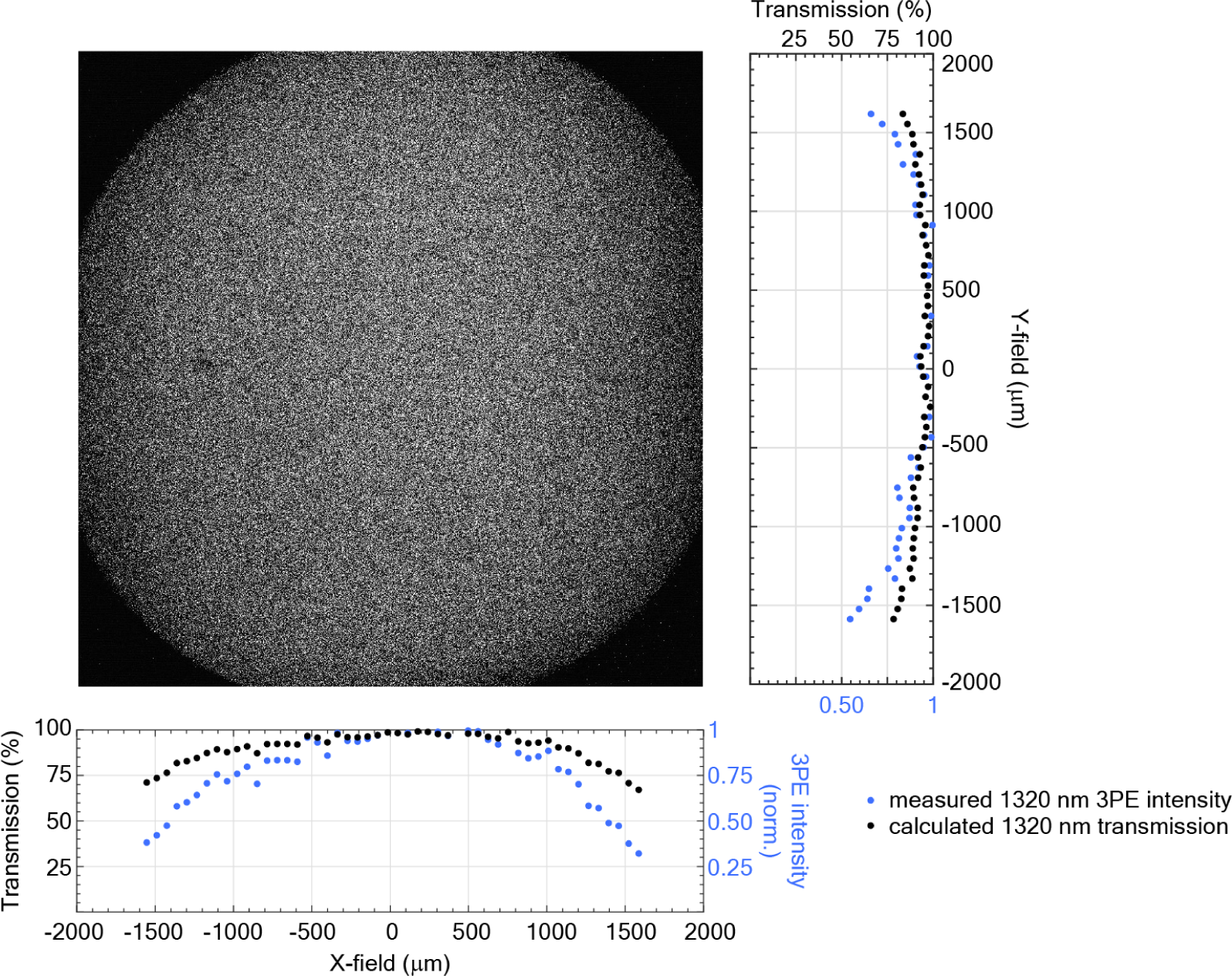
Normalized optical transmission of DEEPscope across the FOV at 1320 nm. Transmission measurement of DEEPscope using a fluorescent dye pool (F36915, Thermo Fisher). The image was acquired by underfilling the polygon scanner and the objective to achieve a uniform axial resolution at ∼20 *μ*m across the entire FOV. The measured X-profile and Y-profile of the image showed the normalized 1320 nm 3P fluorescence intensity across X and Y fields (blue dots). The transmission at 1320 nm (black dots) was calculated by taking a cubic root on the measured data.

**Figure 1—figure supplement 5.**
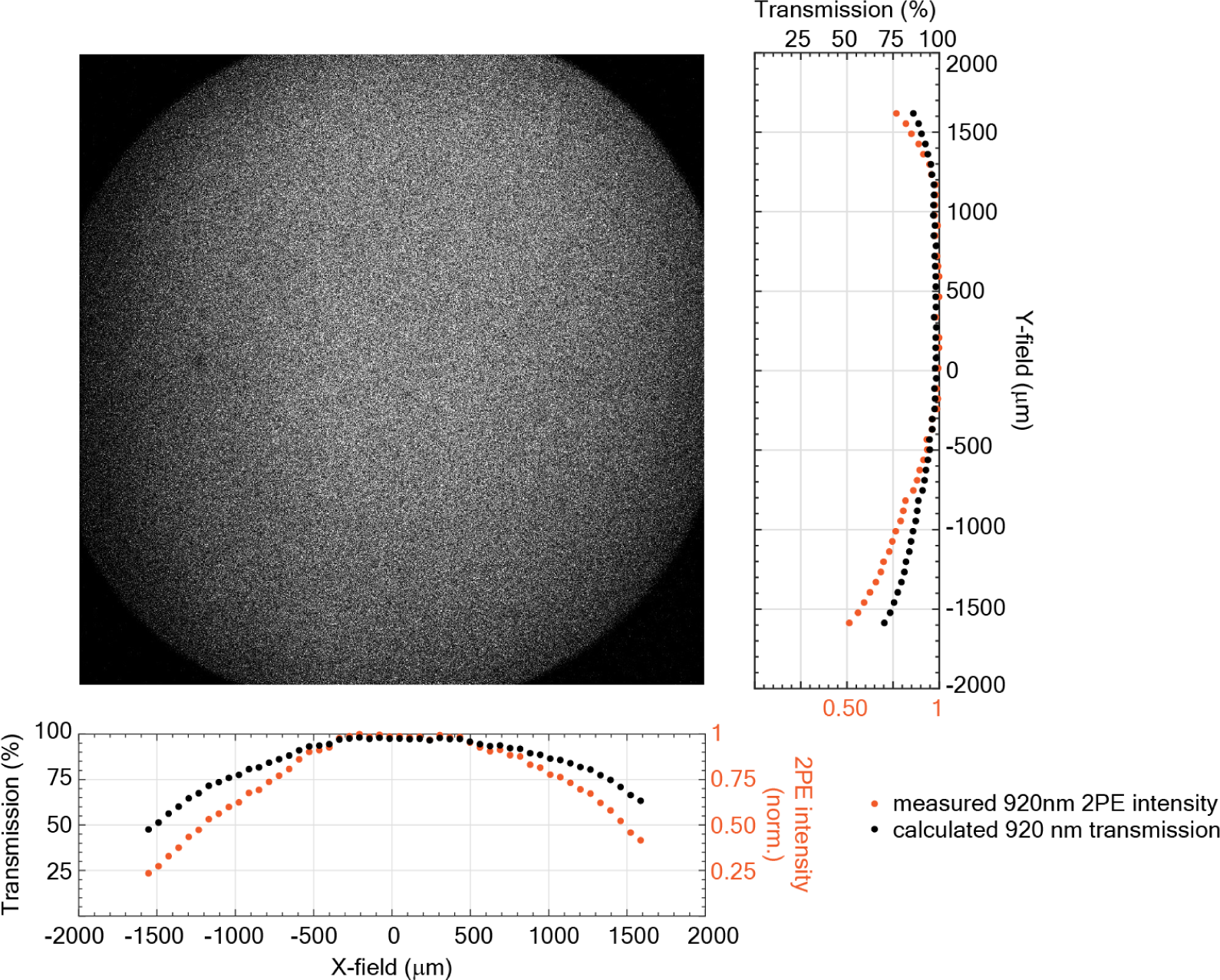
Normalized optical transmission of DEEPscope across the FOV at 920 nm. Transmission measurement of DEEPscope using a fluorescent dye pool (F36915, Thermo Fisher). The image was acquired by underfilling the polygon scanner and objective to achieve a uniform axial resolution at ∼20 um across the entire FOV. The measured X-profile and Y-profile of the image showed the normalized 920 nm 2P fluorescence intensity across X and Y field (orange dots). The transmission at 920 nm (black dots) was calculated by taking a square root on the measured data.

**Figure 1—figure supplement 6.**
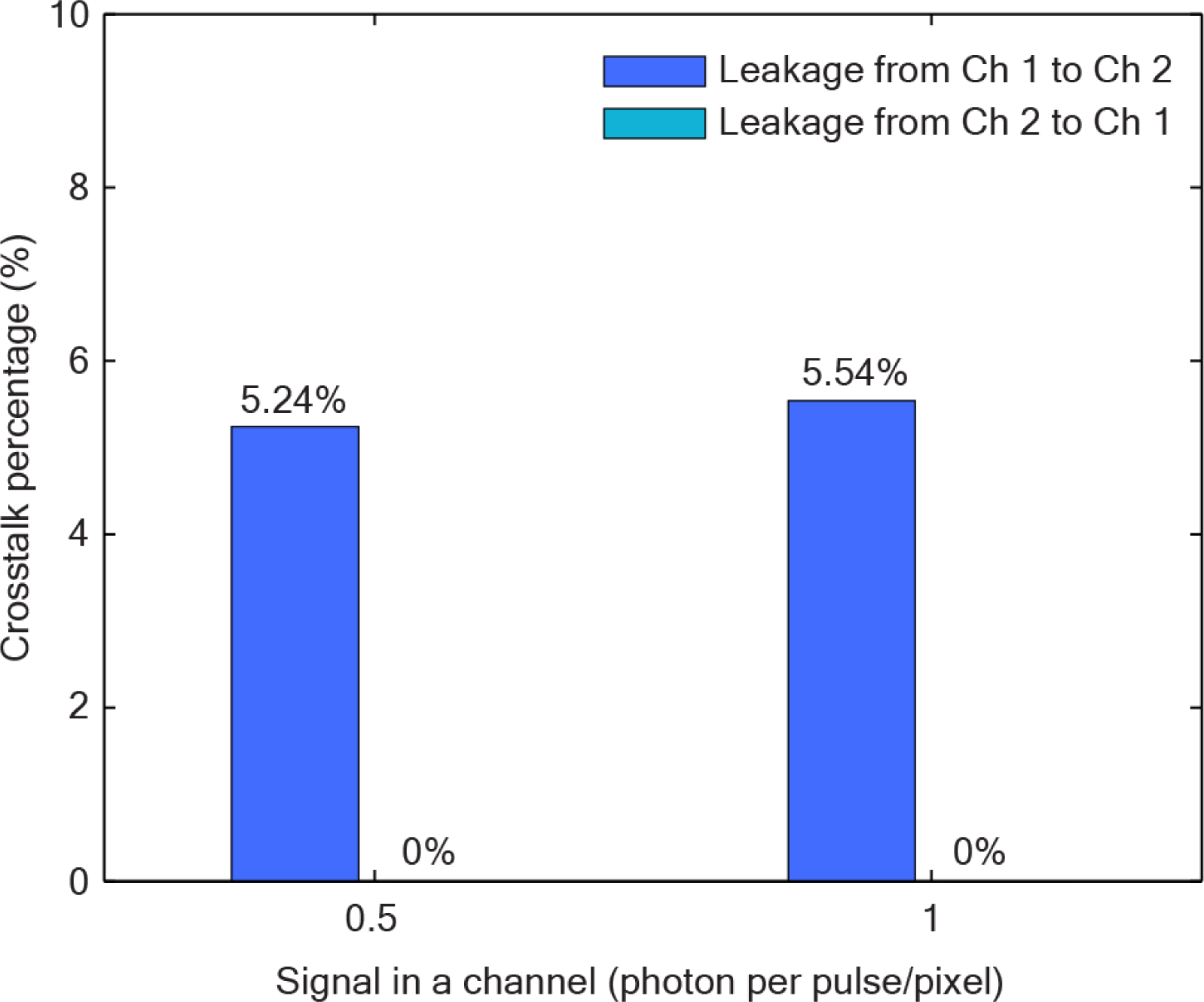
Crosstalk between the fluorescence signal generated by the two beamlets.Measured crosstalk by imaging fluorescein dye pool (F36915, ThermoFisher) with beamlet 1 or 2. The crosstalk percentage from Ch1 to Ch2 is calculated as the mean pixel intensity from Ch 2 divided by the mean pixel intensity from Ch 1 when beamlet 1 is on. The crosstalk percentage from Ch2 to Ch1 was calculated vice versa and is negligible. The PMT gain is ∼10^6^ (0.9V control voltage). This crosstalk was mainly caused by the acquisition bandwidth of the DAQ card of 123 MHz sampling rate and input analog bandwidth of ∼61.5 MHz.

**Figure 1—figure supplement 7.**
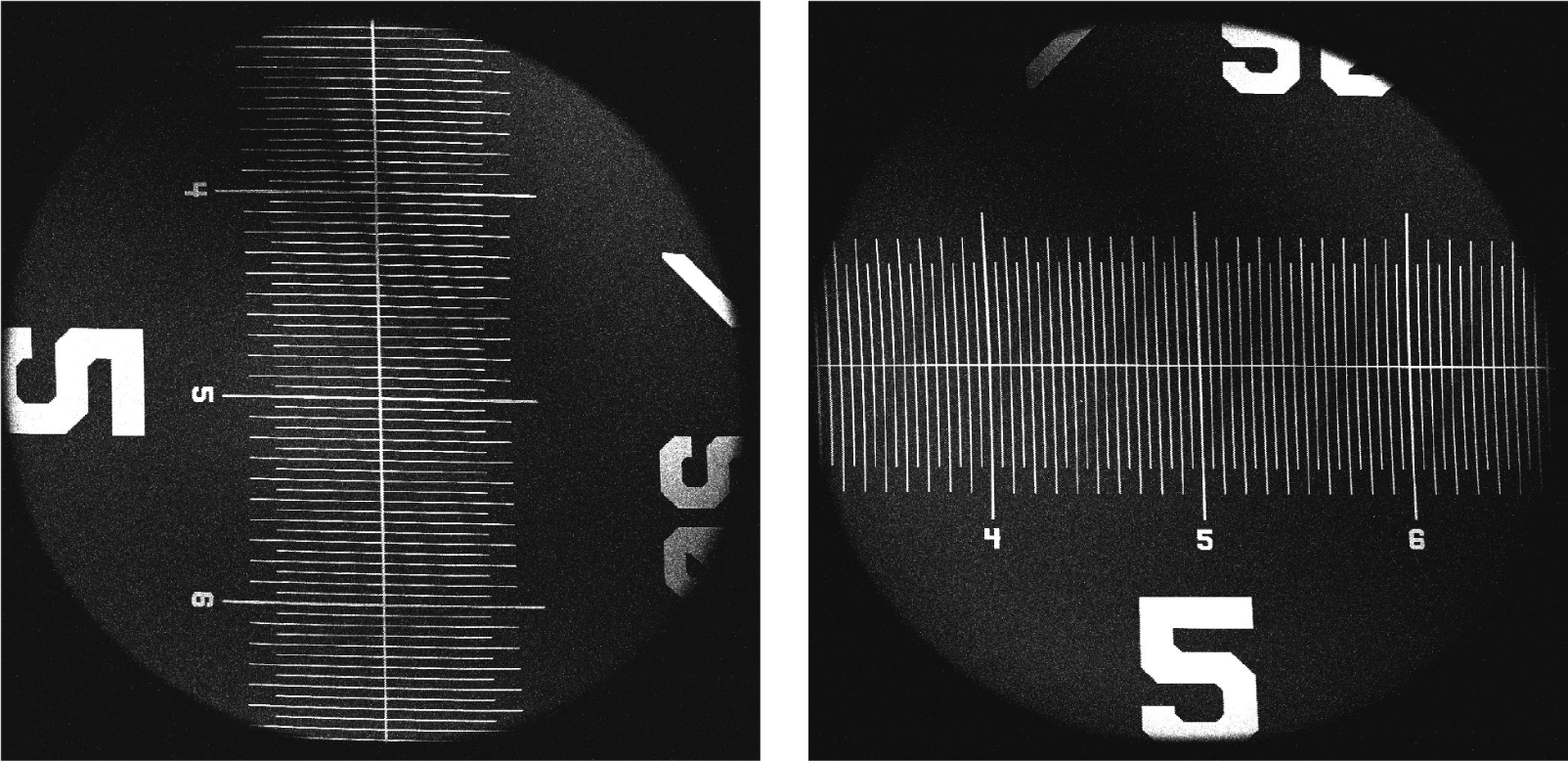
Field-of-view measurement of DEEPscope. THG image of a stage micrometer (R1L3S1P, Thorlabs) obtained by the DEEPscope. It shows a field-of-view of ∼3.5 mm in diameter.

**Figure 1—figure supplement 8.**
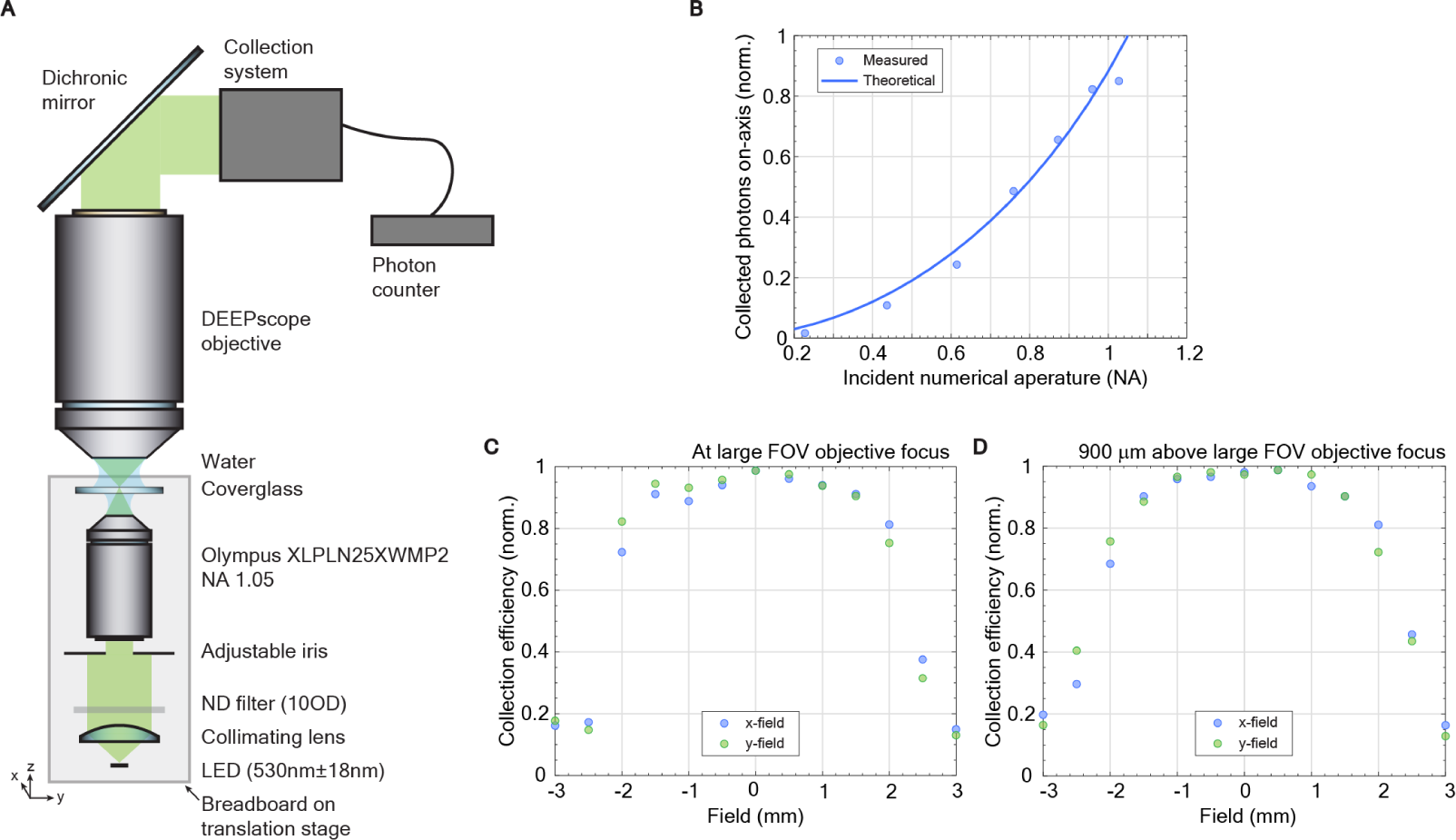
Collection efficiency of the DEEPscope objective lens. **(a.)** Measurement setup for the collection efficiency of DEEPscope objective. **(b.)** Collected photons on-axis at different incident numerical apertures by varying the adjustable iris. It shows that the collected photons start to plateau when the incident numerical aperture is > 1.0, indicating that the collection NA is ∼1.0.**(c.)** Relative collection efficiency at different field positions of the DEEPscope objective at the focus when the incident numerical aperture is ∼1.0.**(d.)** Relative collection efficiency at different field positions of the DEEPscope objective at 900 *μ*m above the focus when the incident numerical aperture is ∼1.0..

**Figure 1—figure supplement 9.**
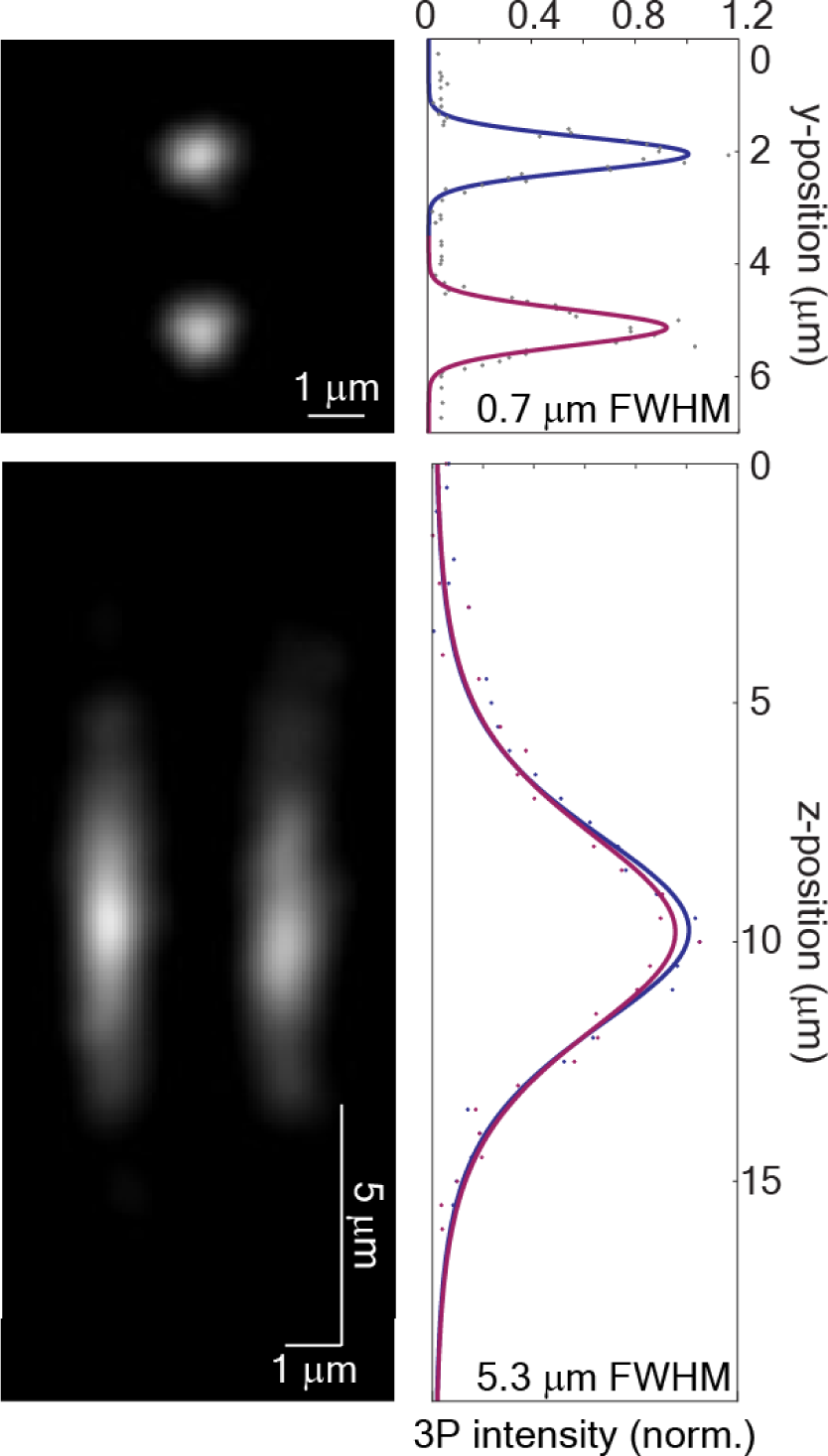
Measured point-spread functions of the two beamlets generated by the beamlet generation delay line. Two co-planar 3P foci with 5.3 **μ**m axial resolution (FWHM) and 0.7 **μ**m lateral resolution (FWHM) were created with the two beamlets at 1320 nm. The foci were ∼3 **μ**m apart along the slow scan axis.

**Figure 1—figure supplement 10.**
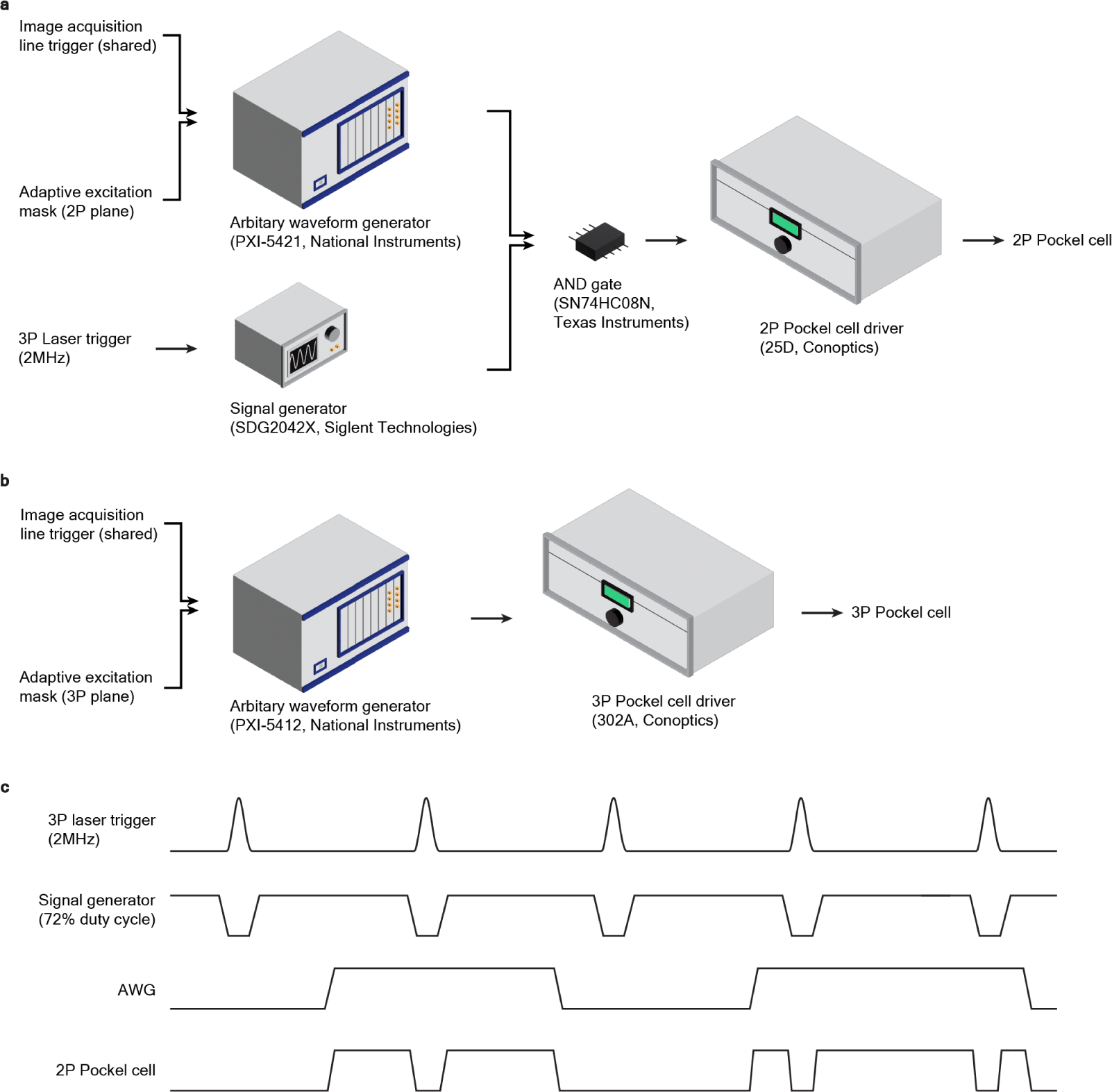
2P/3P multiplexing and dual adaptive excitation.**(a)** Setup for 2P adaptive excitation. An AND gate (SN74HC08N, Texas Instruments) was used to combine 2P/3P multiplexing modulation signal and adaptive excitation modulation**(b)** Setup for 3P adaptive excitation **(c)** Timing diagram for 2P modulation

**Figure 1—figure supplement 11.**
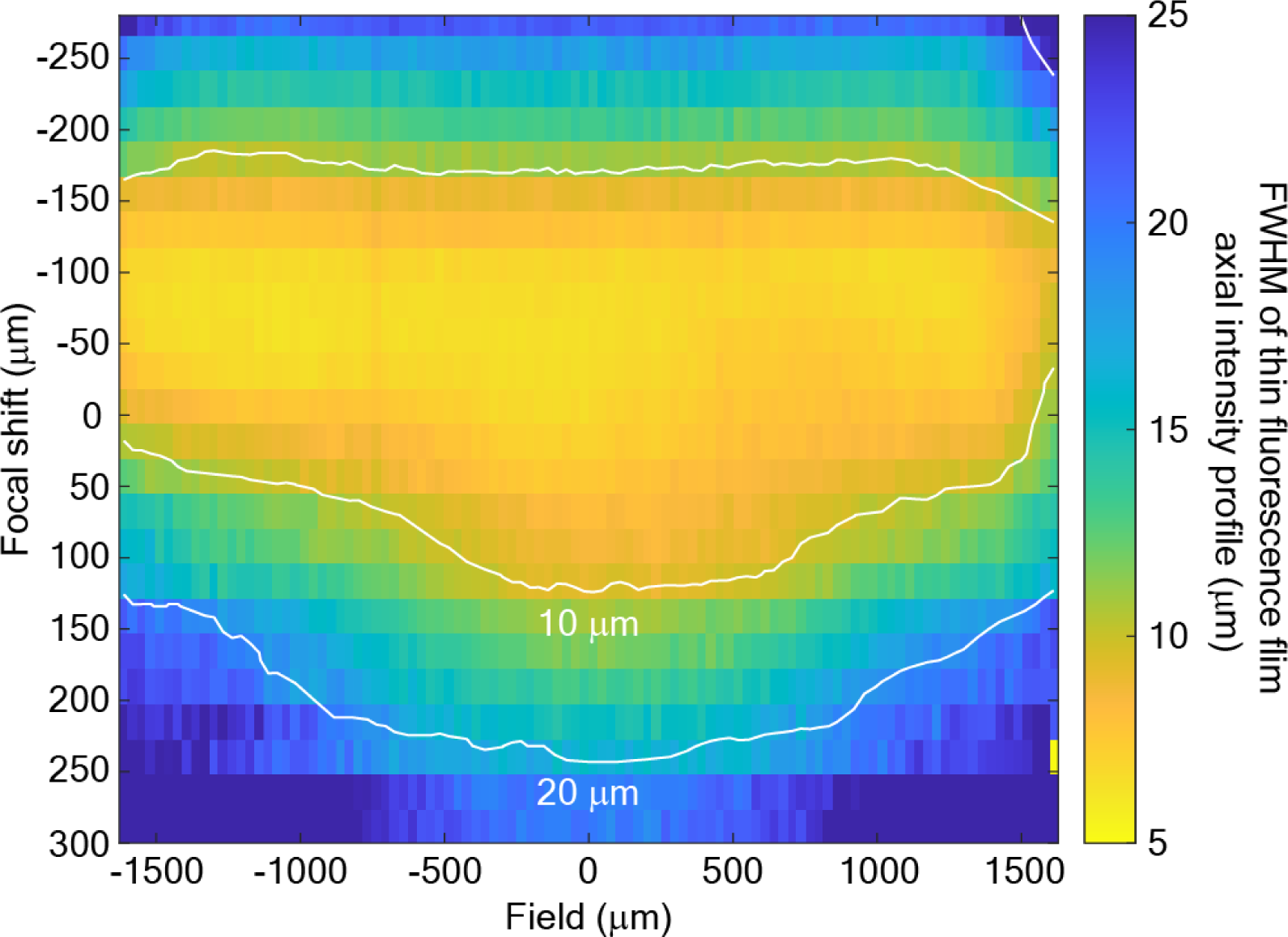
FWHM of the axial intensity profile of a 500-nm thin fluorescent film of Rhodamine B dye with 920 nm two-photon excitation (2PE) across the field-of-view. The data is obtained by axial scanning at different focal positions during remote focusing. X-field and Y-field show similar results.

**Figure 2—figure supplement 1.**
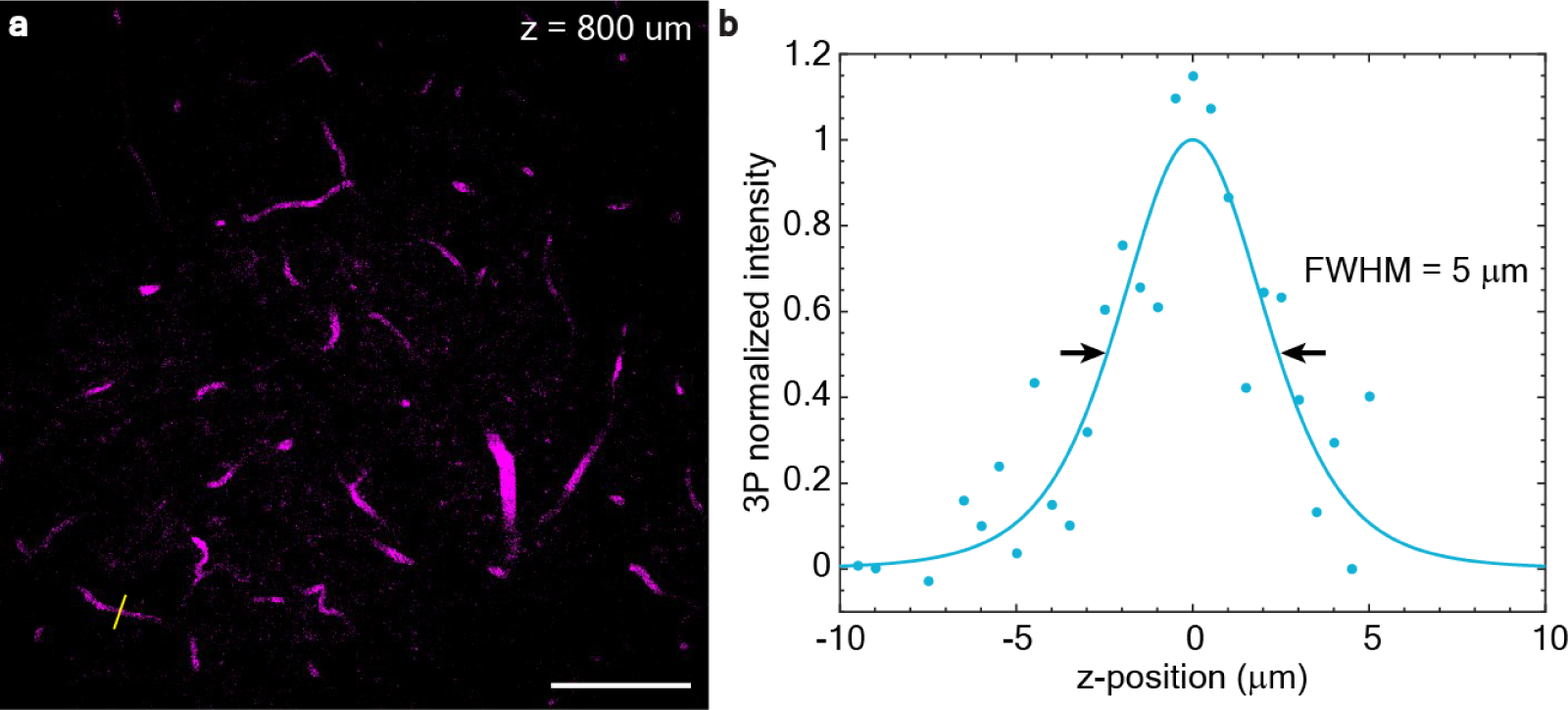
Measurement of 1320 nm 3P axial resolution in-vivo. **(a)** Image of fluorescein-labeled vasculature of a GCaMP6s-expressing transgenic mouse (TIT2L-GC6s-ICL-tTA2, female, 25 weeks) at 800 *μ*m below the brain surface taken with 1320 nm 3P excitation. Scale bar: 100 *μ*m.**(b)** Measurement of axial fluorescence intensity profile across the capillary blood vessel indicated by the yellow line in a, indicating the upper bound of the axial resolution to be ∼ 5 *μ*m.

**Figure 3—figure supplement 1.**
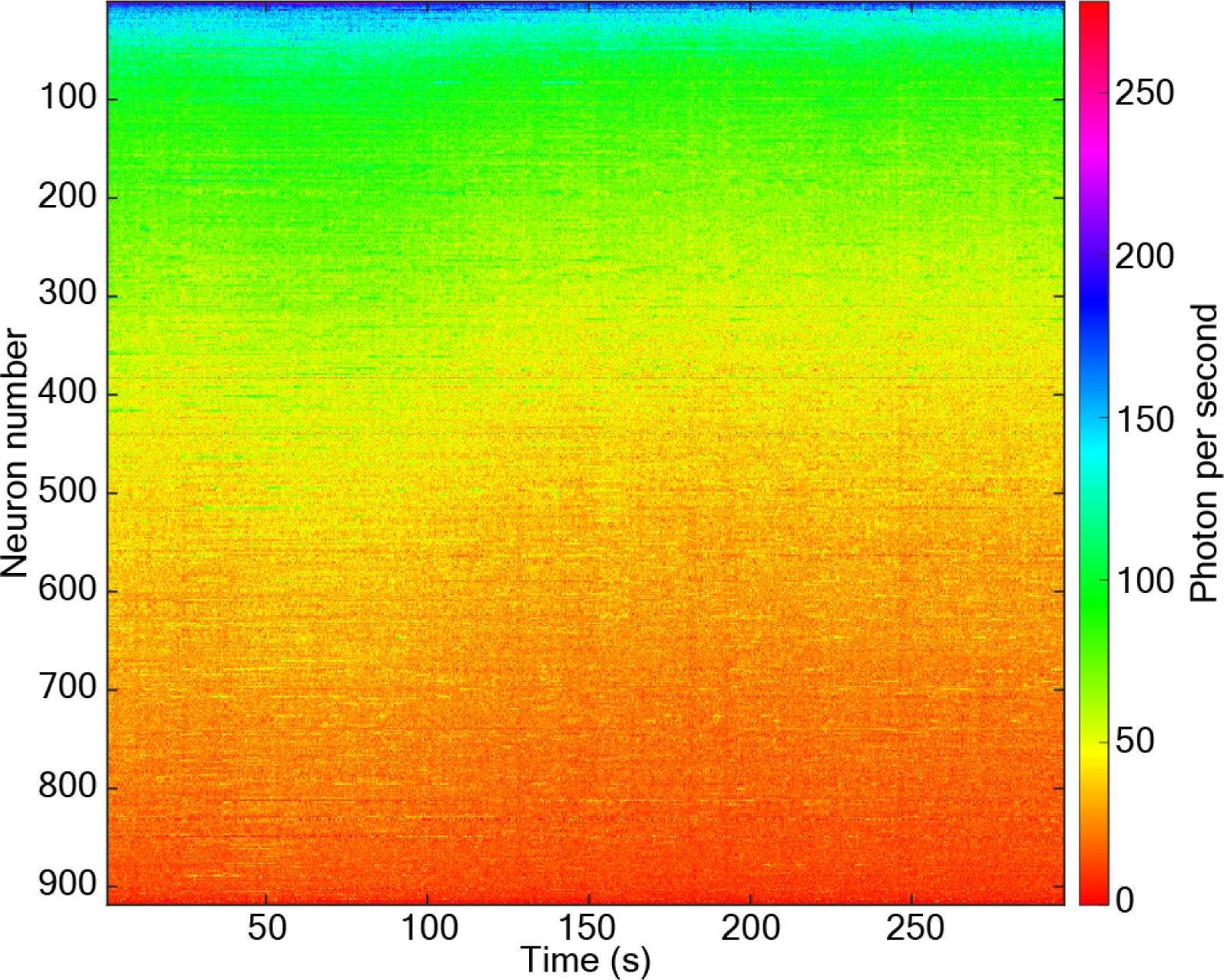
Raw data of neuronal activity traces with 3P DEEPscope. The unprocessed activity traces of neurons in Fig. 3a. The signal of each neuron (in photons/neuron/second) was plotted against time, at the original 4 Hz recording frame rate, without any temporal down-sampling or low-pass filtering.

## References

Beaurepaire E, Mertz J. Epifluorescence collection in two-photon microscopy. Applied optics. 2002; 41(25):5376–5382.

Bounds HA, Sadahiro M, Hendricks WD, Gajowa M, Gopakumar K, Quintana D, Tasic B, Daigle TL, Zeng H, Oldenburg IA, et al. Ultra-precise all-optical manipulation of neural circuits with multifunctional Cre-dependent transgenic mice. bioRxiv. 2021; p. 2021–10.

Chow DM, Sinefeld D, Kolkman KE, Ouzounov DG, Akbari N, Tatarsky R, Bass A, Xu C, Fetcho JR. Deep three-photon imaging of the brain in intact adult zebrafish. Nature Methods. 2020; 17(6):605–608.

Clough M, Chen IA, Park SW, Ahrens AM, Stirman JN, Smith SL, Chen JL. Flexible simultaneous mesoscale two-photon imaging of neural activity at high speeds. Nature Communications. 2021; 12(1):6638.

Dana H, Sun Y, Mohar B, Hulse BK, Kerlin AM, Hasseman JP, Tsegaye G, Tsang A, Wong A, Patel R, et al. High-performance calcium sensors for imaging activity in neuronal populations and microcompartments. Nature methods. 2019; 16(7):649–657.

Demas J, Manley J, Tejera F, Barber K, Kim H, Traub FM, Chen B, Vaziri A. High-speed, cortex-wide volumetric recording of neuroactivity at cellular resolution using light beads microscopy. Nature Methods. 2021; 18(9):1103–1111.

Gauthier JL, Koay SA, Nieh EH, Tank DW, Pillow JW, Charles AS. Detecting and correcting false transients in calcium imaging. Nature methods. 2022; 19(4):470–478.

Janiak F, Bartel P, Bale M, Yoshimatsu T, Komulainen E, Zhou M, Staras K, Prieto-Godino L, Euler T, Maravall M, et al. Non-telecentric two-photon microscopy for 3D random access mesoscale imaging. Nature communications. 2022; 13(1):544.

Li B, Wu C, Wang M, Charan K, Xu C. An adaptive excitation source for high-speed multiphoton microscopy. Nature methods. 2020; 17(2):163–166.

Li Y, Montague SJ, Brüstle A, He X, Gillespie C, Gaus K, Gardiner EE, Lee WM. High contrast imaging and flexible photomanipulation for quantitative in vivo multiphoton imaging with polygon scanning microscope. Journal of Biophotonics. 2018; 11(7):e201700341.

Li Y, Gautam V, Bruestle A, Cockburn I, Daria V, Gillespie C, Gaus K, Alt C, Lee W. Flexible polygon-mirror based laser scanning microscope platform for multiphoton in-vivo imaging. Journal of Biophotonics. 2017; 10(11):1526–1537.

Lu R, Liang Y, Meng G, Zhou P, Svoboda K, Paninski L, Ji N. Rapid mesoscale volumetric imaging of neural activity with synaptic resolution. Nature methods. 2020; 17(3):291–294.

Ota K, Oisi Y, Suzuki T, Ikeda M, Ito Y, Ito T, Uwamori H, Kobayashi K, Kobayashi M, Odagawa M, et al. Fast, cell-resolution, contiguous-wide two-photon imaging to reveal functional network architectures across multi-modal cortical areas. Neuron. 2021; 109(11):1810–1824.

Ouzounov DG, Wang T, Wang M, Feng DD, Horton NG, Cruz-Hernández JC, Cheng YT, Reimer J, Tolias AS, Nishimura N, et al. In vivo three-photon imaging of activity of GCaMP6-labeled neurons deep in intact mouse brain. Nature methods. 2017; 14(4):388–390.

Pachitariu M, Stringer C, Schröder S, Dipoppa M, Rossi LF, Carandini M, Harris KD. Suite2p: beyond 10,000 neurons with standard two-photon microscopy. BioRxiv. 2016; p. 061507.

Rumyantsev OI, Lecoq JA, Hernandez O, Zhang Y, Savall J, Chrapkiewicz R, Li J, Zeng H, Ganguli S, Schnitzer MJ. Fundamental bounds on the fidelity of sensory cortical coding. Nature. 2020; 580(7801):100–105.

Sofroniew NJ, Flickinger D, King J, Svoboda K. A large field of view two-photon mesoscope with subcellular resolution for in vivo imaging. elife. 2016; 5:e14472.

Stirman JN, Smith IT, Kudenov MW, Smith SL. Wide field-of-view, multi-region, two-photon imaging of neuronal activity in the mammalian brain. Nature biotechnology. 2016; 34(8):857–862.

Takasaki K, Abbasi-Asl R, Waters J. Superficial bound of the depth limit of two-photon imaging in mouse brain. Eneuro. 2020; 7(1).

Tsai PS, Mateo C, Field JJ, Schaffer CB, Anderson ME, Kleinfeld D. Ultra-large field-of-view two-photon microscopy. Optics express. 2015; 23(11):13833–13847.

Vladimirov N, Mu Y, Kawashima T, Bennett DV, Yang CT, Looger LL, Keller PJ, Freeman J, Ahrens MB. Light-sheet functional imaging in fictively behaving zebrafish. Nature methods. 2014; 11(9):883–884.

Wang T, Ouzounov DG, Wu C, Horton NG, Zhang B, Wu CH, Zhang Y, Schnitzer MJ, Xu C. Three-photon imaging of mouse brain structure and function through the intact skull. Nature methods. 2018; 15(10):789–792.

Wang T, Wu C, Ouzounov DG, Gu W, Xia F, Kim M, Yang X, Warden MR, Xu C. Quantitative analysis of 1300-nm three-photon calcium imaging in the mouse brain. Elife. 2020; 9:e53205.

Wang T, Xu C. Three-photon neuronal imaging in deep mouse brain. Optica. 2020; 7(8):947–960.

Weisenburger S, Tejera F, Demas J, Chen B, Manley J, Sparks FT, Traub FM, Daigle T, Zeng H, Losonczy A, et al. Volumetric Ca2+ imaging in the mouse brain using hybrid multiplexed sculpted light microscopy. Cell. 2019; 177(4):1050–1066.

Wilt BA, Fitzgerald JE, Schnitzer MJ. Photon shot noise limits on optical detection of neuronal spikes and estimation of spike timing. Biophysical journal. 2013; 104(1):51–62.

Xu C, Webb WW. Multiphoton excitation of molecular fluorophores and nonlinear laser microscopy. In: Topics in Fluorescence Spectroscopy: Volume 5: Nonlinear and Two-Photon-Induced Fluorescence Springer; 2002.p. 471–540.

Yildirim M, Sugihara H, So PT, Sur M. Functional imaging of visual cortical layers and subplate in awake mice with optimized three-photon microscopy. Nature communications. 2019; 10(1):177.

Yu CH, Yu Y, Adsit LM, Chang JT, Barchini J, Moberly AH, Benisty H, Kim J, Young BK, Heng K, et al. The Cousa objective: a long-working distance air objective for multiphoton imaging in vivo. Nature Methods. 2024; 21(1):132–141.

Zhang Y, Rózsa M, Liang Y, Bushey D, Wei Z, Zheng J, Reep D, Broussard GJ, Tsang A, Tsegaye G, et al. Fast and sensitive GCaMP calcium indicators for imaging neural populations. Nature. 2023; 615(7954):884–891.

